# Stacked mutations disrupting syringyl and *p*-coumaroylated lignin biosynthesis in rice result in lignin dominated by guaiacyl units: insights into grass-specific lignin monomer biosynthesis and polymerization mechanisms

**DOI:** 10.1101/2025.03.23.644785

**Authors:** Pingping Ji, Osama A. Afifi, Senri Yamamoto, Yuriko Osakabe, Keishi Osakabe, Toshiaki Umezawa, Yuki Tobimatsu

## Abstract

- The aromatic composition of lignin significantly impacts the usability of lignocellulosic biomass. In eudicots, transgenic and mutant lines with elevated guaiacyl (G) or syringyl (S) lignin units have been successfully generated by manipulating the expression level of CONIFERALDEHYDE 5-HYDROXYLASE (CAld5H). However, this bioengineering approach has proven less effective in grasses, implicating the potential existence of a grass-specific alternative pathway for S lignin biosynthesis.
- Through characterization of genome-edited rice mutants, we demonstrated that S lignin in rice can be virtually eliminated by disrupting genes encoding CAld5H along with *p*- COUMAROYL-COENZYME A:MONOLIGNOL TRANSFERASE (PMT), a grass-specific enzyme essential for the biosynthesis of monolignol *p*-coumarate conjugates. In contrast, individual mutations in either *CAld5H* or *PMT* genes resulted in incomplete elimination of S lignin. These findings provide strong evidence that rice possesses a CAld5H-independent pathway leading to the grass-specific monolignol *p*-coumarate conjugates.
- In-depth structural characterizations of G-dominated lignins from rice and Arabidopsis mutants, natural gymnosperm pine, and G-type synthetic lignin revealed pronounced effects of lineage-dependent cell wall environments on the linkage patterns and molecular weight distributions of the resulting lignin polymers.
- Overall, our findings highlight previously overlooked lineage-specific lignin monomer biosynthesis and polymerization patterns in grasses.

## Introduction

Grass lignocellulose, along with wood lignocellulose, represents a sustainable resource for producing biomass-based biochemicals and biofuels through biorefining. Grass biomass crops, such as sugarcane (*Saccharum* spp.), sorghum (*Sorghum bicolor*), switchgrass (*Panicum virgatum*), and *Miscanthus* spp., are particularly valued for their high lignocellulose productivity (Mullet *et al*., 2014; Tye *et al*., 2016; Umezawa *et al*., 2020). Additionally, agricultural residues from grass grain and sugar crops offer significant potentials in utilizing them in biorefinery (Lal, 2005). Compared to woody biomass, grass lignocellulosic biomass is also more amenable to chemical delignification, facilitating isolations of polysaccharide components for further downstream product development (Davis *et al*., 2013; Tye *et al*., 2016; Bhatia *et al*., 2017; Umezawa *et al*., 2020). However, as discussed further below, the diverse and complex nature of grass lignocellulose components, particularly lignin, presents challenges in biorefinery. This underscores the importance of understanding the biosynthesis and structural characteristics of grass lignocellulose to optimize its industrial applications through molecular breeding and bioengineering approaches (Barros and Dixon, 2020; Umezawa *et al*., 2020; Chandrakanth *et al*., 2023; Peracchi *et al*., 2024; Umezawa, 2024).

Lignin is an aromatic biopolymer formed through the oxidative radical coupling, or dehydrogenative polymerization, of *p*-hydroxycinnamyl alcohols (monolignols) and related compounds in the cell walls (Freudenberg and Neish, 1968; Sarkanen and Ludwig, 1971; Adler, 1977; Higuchi, 1985; Ralph *et al*., 2004, 2009, 2019). Lignin substructure exhibits significant variability across plant lineages, primarily due to the diversity of lignin monomers involved in lignification (Ralph *et al*., 2019). Generally, ferns and gymnosperms produce lignins predominantly composed of guaiacyl (G) units derived from G-type monolignol (coniferyl alcohol). On the other hand, eudicots among angiosperms incorporate both syringyl (S) units derived from S-type monolignol (sinapyl alcohol) and G units. Both gymnosperm and eudicot lignins also contain smaller amounts of *p*-hydroxyphenyl (H) units from H-type monolignol (*p*-coumaryl alcohol). Monocotyledonous grasses among angiosperms, however, exhibit a further unique and complex lignin composition. In addition to G, S, and H units derived from the conventional (non-acylated) monolignols, grass lignins incorporate monolignol *p*-coumarate conjugates, such as γ-*p*-coumaroylated G- and S-type monolignols (coniferyl and sinapyl *p*-coumarate), which give rise to G and S units, respectively, appended with grass-specific *p*-coumarate (*p*CA) units. Furthermore, grass lignins incorporate feruloylated monolignols and arabinoxylan, and the flavonoid tricin, leading to the formation of grass-specific ferulate (FA) and tricin units in the final lignin polymer (**Fig. 1**) (Ralph *et al*., 2019).

**Fig. 1.**
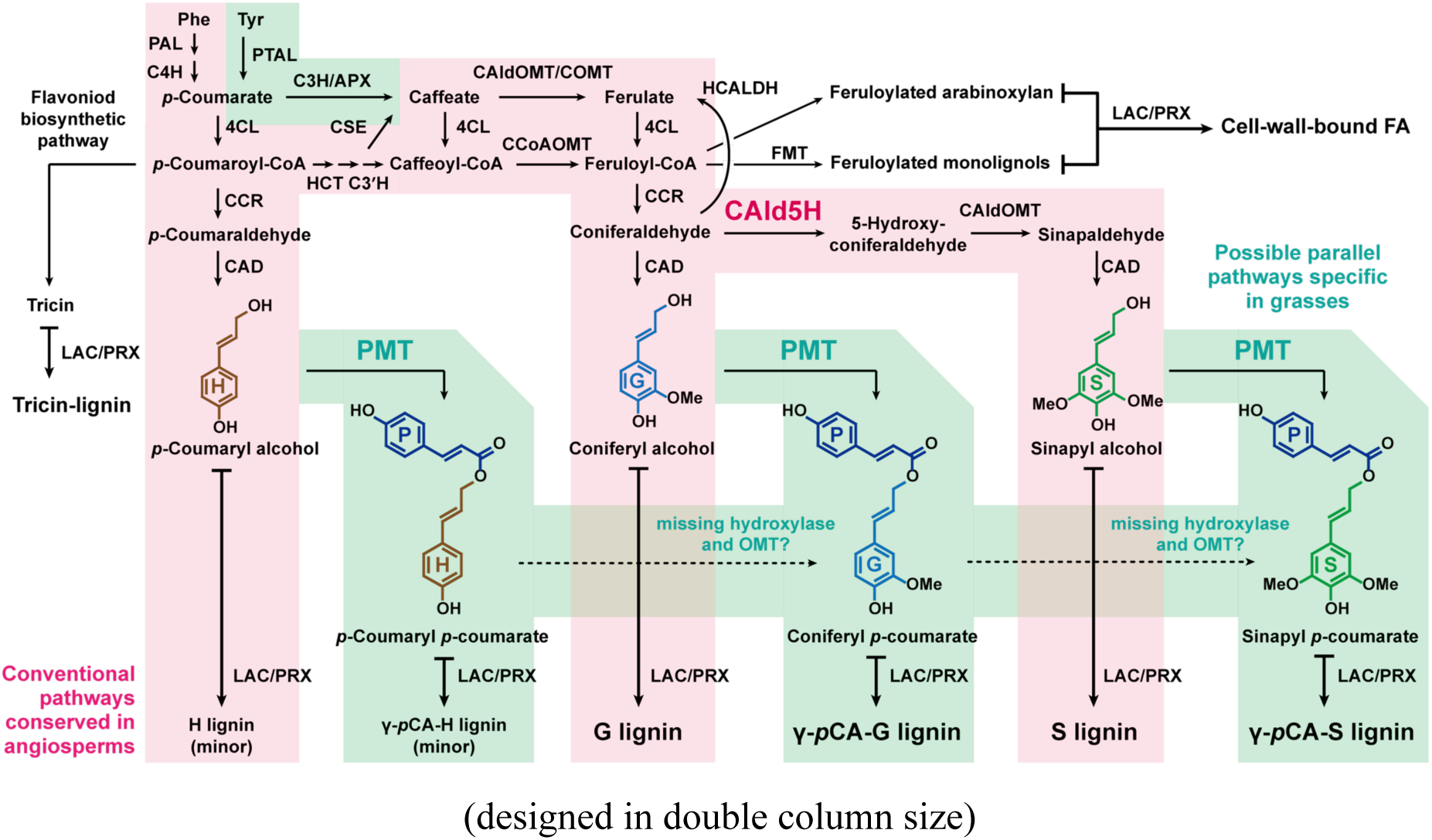
Proposed lignin biosynthetic pathways in grasses. PAL, phenylalanine ammonia-lyase; PTAL, phenylalanine/tyrosine ammonia-lyase; C4H, cinnamate 4-hydroxylase; C3H/APX, 4-coumarate 3-hydroxylase/ascorbate peroxidase; 4CL, 4-coumarate:CoA ligase; C3’H, *p*-coumaroyl ester 3-hydroxylase; HCT, *p*-hydroxycinnamoyl-CoA:quinate/shikimate transferase; CSE, caffeoyl shikimate esterase; CCoAOMT, caffeoyl-CoA *O*-methyltransferase; CAldOMT, 5-hydroxyconiferaldehyde *O*-methyltransferase; COMT, caffeate *O*-methyltransferase; CCR, cinnamoyl-CoA reductase; HCALDH, hydroxycinnamaldehyde dehydrogenase; CAld5H, coniferaldehyde 5-hydroxylase; CAD, cinnamyl alcohol dehydrogenase; PMT, *p*-coumaroyl-CoA:monolignol transferase; FMT, feruroyl-CoA:monolignol transferase; LAC, laccase; PRX, peroxidase; H, *p*-hydroxyphenyl units; G, guaiacyl units; S, syringyl units; P, *p*-coumarate units. The CAld5H (magenta) and PMT (green) of interest in this study, and the conventional monolignol biosynthetic pathways conserved in angisperms (magenta) and possible parallel monolignol *p*-coumarate pathways specific in grasses (green) are highlighted.

Due to the significant impact of lignin aromatic composition on lignocellulosic biomass utilization, numerous efforts have been dedicated to genetically engineering lignin aromatic composition in both eudicot and grass model plants. CONIFERALDEHYDE 5-HYDROXYLASE (CAld5H or FERULATE 5-HYDROXYLASE, F5H), a cytochrome P450 (CYP) enzyme belonging to the CYP84 family, catalyzes the 5-hydroxylation step that converts G-type monolignol precursors into S-type monolignol precursors (Humphreys *et al*., 1999; Osakabe *et al*., 1999) (**Fig. 1**). This enzyme has been shown to effectively modulate the ratio of S and G lignin units, the two major aromatic types of angiosperm lignins, in several eudicot model species. For instance, down-regulation of *CAld5H* genes results in considerably decreased or even absent S lignin content (Meyer *et al*., 1998; Marita *et al*., 1999; Reddy *et al*., 2005; Anderson *et al*., 2015). Conversely, over-expression of *CAld5H* genes increases S lignin content and in a severe case can result in an excess of 90% S lignin (Humphreys *et al*., 1999; Franke *et al*., 2000; Li *et al*., 2003; Stewart *et al*., 2009; Anderson *et al*., 2015). Studies also demonstrated that heterologous expression of eudicot *CAld5H* genes in gymnosperms, which typically produce only G lignins in the cell walls, leads to ectopic productions of S lignin in the walls (Wagner *et al*. 2015; Edmunds *et al*., 2017). Overall, these data strongly support the indispensable role of CAld5H in S lignin biosynthesis in eudicots.

While the conservation of CAld5H in monocot grass lignin biosynthesis has been well-confirmed (Bewg *et al*., 2016; Takeda *et al*., 2017, 2019a; Wu *et al*., 2019; Tetreault *et al*., 2020; Shafiei *et al*., 2023), however, the capacity to modulate the S/G lignin unit ratio in grasses by manipulating CAld5H expression appears to be less straightforward and effective compared to eudicots. In rice, a loss-of-function of *OsCAld5H1*, a primary *CAld5H* gene in rice, resulted in a predictable enrichment of G lignin units but only a partial reduction of S lignin units in major vegetative tissues. Detailed structural analyses of altered lignins produced by the *CAld5H*-deficient rice mutants revealed that enrichment of G units was limited to the non-γ-*p*-coumaroylated units, leaving grass-specific γ-*p*-coumaroylated lignin units largely unaffected (Takeda *et al*., 2019a). These observations, together with supporting data from other grass mutant and transgenic lines (Barros *et al*., 2016; Koshiba *et al*., 2017; Takeda *et al*., 2018; Miyamoto *et al*., 2019; Miyamoto *et al*., 2020; Afifi *et al*., 2022; Takeda-Kimura *et al*., 2025), led to the notion that rice and possibly other grasses may possess parallel monolignol pathways independent of CAld5H activity to produce the grass-specific monolignol *p*-coumarate conjugates separate from the conventional non-γ-*p*-coumaroylated monolignols (Takeda *et al*., 2019a; Barros and Dixon, 2020; Umezawa *et al*., 2020; Peracchi *et al*., 2024; Umezawa, 2024) (**Fig. 1**). However, due to a limited understanding of these potential parallel pathways, effective regulation of the S/G lignin unit ratio in grasses has not yet been fully addressed.

In this study, we aimed to develop rice mutants enriched in G lignin units by introducing stacked mutations that block both conserved and grass-specific S-type lignin monomer biosynthesis. Based on the notion that rice possesses a *CAld5H*-independent pathway leading to monolignol *p*-coumarate conjugates, we introduced loss-of-function mutations to *OsCAld5H1* using CRISPR-Cas9-mediated targeted mutagenesis in a rice mutant lacking *p*-COUMAROYL-COENZYME A:MONOLIGNOL ACYLTRANSFERASE (PMT), a grass-specific enzyme essential for generating monolignol *p*-coumarate conjugates (Withers *et al*., 2012; Lam *et al*., 2024) (**Fig. 1**). As anticipated, we successfully obtained rice mutants with lignins dominated by G units and lacking S units. The structural features of the G-dominated lignins produced by these rice mutants were characterized using a comprehensive suite of analytical techniques, including chemical methods, two-dimensional (2D) nuclear magnetic resonance (NMR), and gel permeation chromatography (GPC). These analyses were conducted comparatively with other G-dominated lignins from a eudicot *CAld5H*-deficient Arabidopsis mutant and natural gymnosperm pine, as well as synthetic lignin polymer (dehydrogenation polymer, DHP) prepared by *in vitro* polymerization of G-type monolignol (coniferyl alcohol). Taken together, our findings provide further evidence for the presence of the grass-specific parallel monolignol pathway and insights into the influence of the lineage-specific cell wall environment on lignin polymerization.

## Materials and Methods

### Plant materials

*oscald5h1-1* and *oscald5h1-2* (identical to *OsCAld5H1*-KO-a-9 and *OsCAld5H1*-KO-c-1, respectively, reported in Takeda *et al*., 2019a) and *ospmt1 ospmt2* (identical to *ospmt1 ospmt2-2* reported in Lam *et al*., 2024) rice mutants were obtained through CRISPR-Cas9-mediated genome editing of rice (cv. Nipponbare) as described earlier (Toda *et al*., 2019; Yamamoto *et al*., 2024). To generate *oscald5h1-3 ospmt1 ospmt2* and *oscald5h1-4 ospmt1 ospmt2*, two single-guide RNAs (sgRNA1 and sgRNA2) targeting the first exon of *OsCAld5H1* (LOC_Os10g36848) (**Fig. 2a**) were designed using the CRISPR-P 2.0 program (Liu *et al*., 2017) and integrated into the multiplex CRISPR-Cas9 binary vector pMgPoef4_129-2A-GFP (Toda *et al*., 2019; Yamamoto et al., 2024) using the Golden Gate Assembly Mix (New England Biolabs, USA) and the oligonucleotides listed in **Table S1**. The binary vector was then transformed into embryogenic calli from *ospmt1 ospmt2* via *Agrobacterium tumefaciens* stain EHA101 and regenerated following the methods of Hiei *et al*. (1994). The regenerated T_0_ plants were genotyped and grown to maturity in potting soil in a growth chamber (Lam *et al*., 2017). Selected T_1_ plants were further genotyped and grown to maturity in a greenhouse maintained at 27 ℃ (Lam *et al*., 2017) along with the wild-type rice and *oscald5h1-1*, *oscald5h1-2*, and *ospmt1 ospmt2* mutant lines. For genotyping, genomic DNA was extracted from young leaves, and the genomic region containing the target site was amplified by PCR using primers listed in **Table S1**. The amplified PCR products were subjected to direct sequencing according to the method of Takeda *et al*. (2019a). Arabidopsis *atcald5h1* (*fah1-2*; accession: CS6172) mutant (Meyer *et al*., 1998) was obtained from the Arabidopsis Biological Resource Center at Ohio University, USA, and grown to maturity in an incubator maintained at 22 °C with a 16-hour light/8-hour dark photoperiod, along with Arabidopsis Col-0 ecotype used as a wild-type control. The origin of mature *Pinus taeda* wood sample used in this study is detailed in Miyagawa *et al*. (2020).

**Fig. 2.**
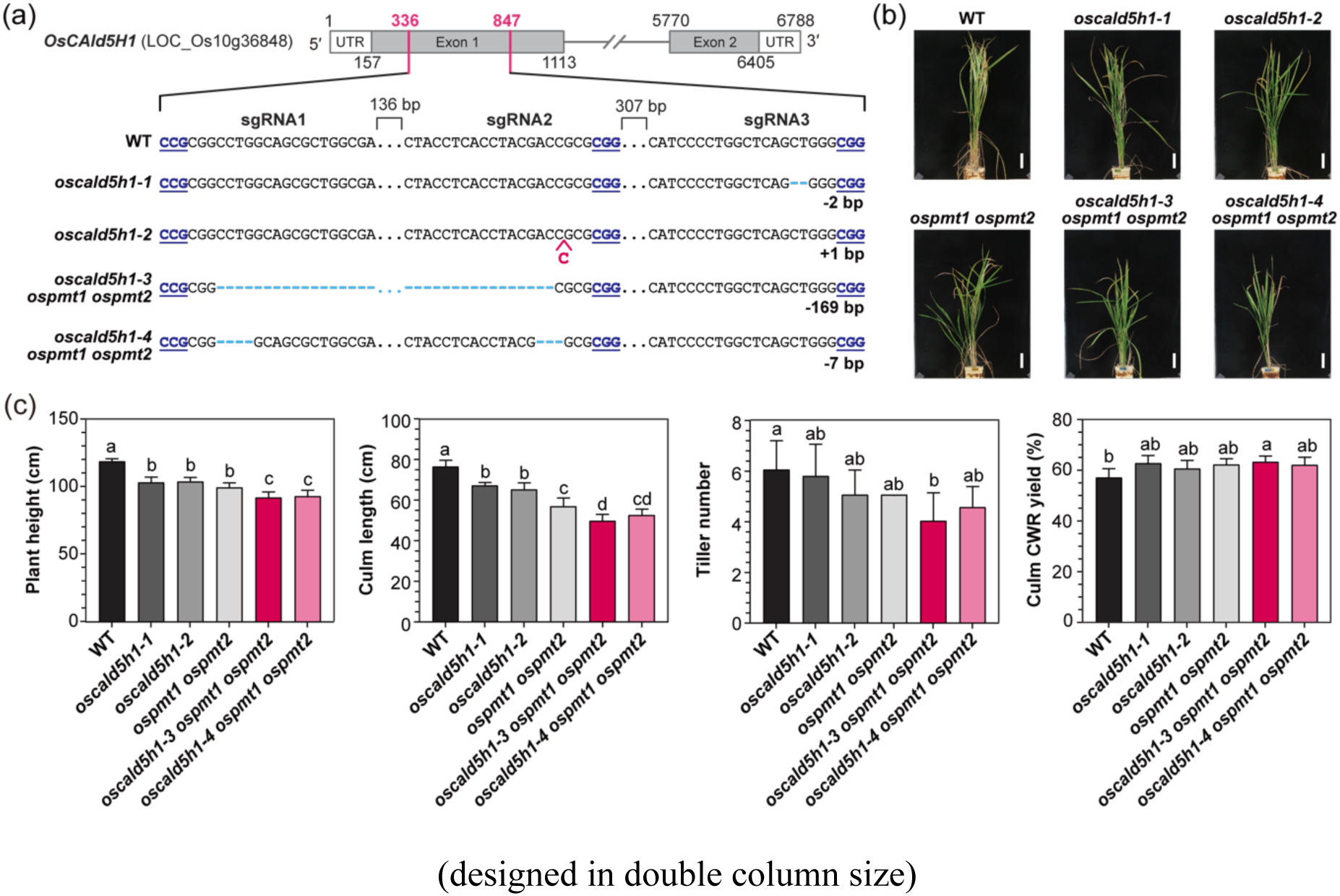
Generation of *oscald5h1 ospmt1 ospmt2* rice mutants. (a) Gene structure and mutation patterns in the *OsCAld5H1* (LOC_Os10g36848) locus of *oscald5h1* (Takeda et al., 2019) and *oscald5h1 ospmt1 ospmt2* (this study) mutants generated by CRISPR-Cas9-mediated targeted mutagenesis. The protospacer adjacent motif (PAM; underlined), inserted (colored in red) and deleted (colored in blue) sequences are highlighted. sgRNA1 and sgRNA2 were used to generate *oscald5h1-3 ospmt1 ospmt2* and *oscald5h1-4 ospmt1 ospmt2* mutants using *ospmt1 ospmt2* (Lam et al., 2024) as a background in this study. sgRNA3 was used to generate *oscald5h1-1* and *oscald5h1-2* mutants using wild-type rice as a background previously (Takeda et al., 2019). (b and c) Morphological appearance (b) and growth parameters (c) of mature wild-type and mutant (T1 homozygous lines) rice plants grown under greenhouse conditions. Scale bar = 10 cm. Values in (c) are means ± standard deviation from biological replicates (*n* = 4–5). Different letters on bars indicate significant differences (one-way ANOVA with Tukey’s test, P < 0.05). WT, wild-type line; *oscald5h1-1* and *oscald5h1-2*, *OsCAld5H1* single-knockout lines; *ospmt1 ospmt2*, *OsPMT1* and *OsPMT2* double-knockout lines; *oscald5h1-3 ospmt1 ospmt2* and *oscald5h1-4 ospmt1 ospmt2*, *OsCAld5H1*, *OsPMT1* and *OsPMT2* triple-knockout lines.

### Cell wall and lignin sample preparations

Dried mature rice culms, Arabidopsis stems, and *P. taeda* wood were pulverized using a TissueLyser II (Qiagen, Hilden, Germany) shaker mill, and then sequentially extracted with water, 80% aqueous ethanol and acetone, to prepare the cell wall residue (CWR) samples used for chemical and NMR analyses (Mansfield *et al*., 2012). The dioxane/water-soluble lignin (DL) samples used for NMR and GPC were prepared according to the method described previously (Martin *et al*., 2023; Lam *et al*., 2024). Briefly, CWR samples (approximately 1g pooled from more than 4 and 20 plants for rice and Arabidopsis, respectively) were further ball-milled using a Pulverisette7 (Fritsch Industrialist, Idar-Oberstein, Germany) planetary ball-mill in zirconium vessels containing zirconium ball bearings (600 rpm, 24 cycles of 10 min milling and 5 min break) and then subjected to digestion with crude cellulases (CELLULYSIN^®^, Merk Millipore, Darmstadt, Germany). The digested CWR samples were then extracted with dioxane-water (96:4, v/v) and purified by reprecipitation in 0.01 M hydrochloric acid aqueous solution. G-type synthetic lignin (G-DHP) was prepared by *in vitro* polymerization of coniferyl alcohol using the end-wise polymerization method according to the previously described method (Tobimatsu *et al*., 2013).

### Chemical analysis

The Klason lignin assay (Hatfield *et al*., 1994), quantification of cell wall-bound FA and *p*CA by mild alkaline hydrolysis (Yamamura *et al*., 2011), neutral sugar analysis by two-step trifluoroacetic acid/sulfuric acid-catalyzed hydrolysis (Lam *et al*., 2017), and derivatization followed by reductive cleavage (DFRC) (Karlen *et al*., 2018; Takeda *et al*., 2018) were conducted according to the methods described previously.

### 2D NMR analysis

For the 2D NMR analysis of the whole cell walls from rice stems, CWR samples (pooled from 4 biologically independent plants) were finely ball-milled using a Pulverisette7 (Fritsch Industrialist, Idar-Oberstein, Germany) planetary ball-mill following previously described conditions (Kim and Ralph, 2010). For the whole cell wall NMR analysis, aliquots of the ball-milled CWR samples (approximately 60 mg) were swelled in 600 μL dimethylsulfoxide-*d*_6_/pyridine-*d*_5_ (4:1, v/v) and then subjected to the ^1^H–^13^C heteronuclear single-quantum coherence (HSQC) NMR analysis (Kim and Ralph, 2010; Mansfield *et al*., 2012). For the NMR analysis of DL and G-DHP samples, the samples (approximately 20 mg) were dissolved in 500 μL dimethylsulfoxide-*d*_6_/pyridine-*d*_5_ (4:1, v/v) and then subjected to the 2D HSQC NMR analysis (Martin *et al*., 2023; Lam *et al*., 2024). The HSQC NMR spectra were acquired using a Bruker Avance III 800 system (800 MHz; Bruker Biospin, Billerica, MA, USA) equipped with a cryogenically cooled 5 mm TCI gradient probe.

Adiabatic HSQC experiments were conducted using the standard Bruker implementation (‘hsqcetgpsp.3’) and previously described parameters (Kim and Ralph, 2010; Mansfield *et al*., 2012). Data were processed and analyzed using the TopSpin 4.3 software (Bruker Biospin) as previously described (Afifi *et al*., 2022; Martin *et al*., 2023; Lam *et al*., 2024). Peak assignment was based on comparison with NMR data in literature (Kim and Ralph, 2010; Mansfield *et al*., 2012; Lan *et al*., 2015; 2018; Afifi *et al*., 2022; Martin *et al*., 2023; Lam *et al*., 2024; Ralph *et al*., 2024). The central dimethylsulfoxide-*d*_6_ solvent peaks (δ_C_/δ_H_, 39.5/2.49 ppm) were used for calibrating the chemical shift.

### GPC

The GPC analysis of DL and G-DHP samples was conducted following the method described previously (Tobimatsu *et al*., 2013). Briefly, the lignin samples were dissolved in *N,N*-dimethylformamide with 0.1 M lithium bromide (approximately 3 mg mL^−1^), and subjected to GPC analysis on a Shimadzu LC-20AD LC system equipped with SPD-20A UV/VIS detector using the following conditions: column: Tosoh TSK gel α-M + α-2500; eluent: *N,N*-dimethylformamide with 0.1 M lithium bromide; flow rate: 0.5 mL min^−1^; column oven temperature: 40 °C; sample detection: UV abdorption at 280 nm. The data acquisition and computation were performed using a Shimadzu LCsolution version 5.9 software. Molecular weight calibration was conducted using the Agilent EasyVials polystyrene standards (Agilent Technologies, Santa Clara, CA, USA).

### Statistical analysis

Student’s *t* test and one-way ANOVA followed by Tukey’s multiple comparison test were performed using the GraphPad Prism version 8 (GraphPad Software, San Diego, CA, USA).

### Accession numbers

The accession numbers for *CAld5H* and *PMT* genes studied in this study are listed in **Table S2**.

## Results

### Generation of *oscald5h1 ospmt1 ospmt2* rice mutants

To generated rice mutants with stacked mutations in both *CAld5H* and *PMT* genes, we introduced additional *OsCAld5H1* mutations into the *ospmt1 ospmt2* line, a genome-edited mutant line carrying loss-of-function mutations in *OsPMT1* and *OsPMT2*, two primary genes encoding PMT in rice (Lam *et al*., 2024). As a result, we successfully isolated two homozygous triple-knockout mutant lines, *oscald5h1-3 ospmt1 ospmt2* and *oscald5h1-4 ospmt1 ospmt2*, which harbor −169 bp and −7 bp deletions in the targeted *OsCAld5H1* site (**Fig. 2a**).

Fully genotyped *oscald5h1-3 ospmt1 ospmt2* and *oscald5h1-4 ospmt1 ospmt2* mutants (T_1_ progenies) were grown alongside with wild-type control (WT), as well as the previously generated *CAld5H*-deficient *oscald5h1-1* and *oscald5h1-2* (**Fig. 2a**) (Takeda *et al*., 2019a) and *PMT*-deficient *ospmt1 ospmt2* (Lam *et al*., 2024) mutant lines for phenotype and cell wall characterizations (**Fig. 2b**). As previously reported (Takeda *et al*., 2019a; Lam *et al*., 2024), *oscald5h1* and *ospmt1 ospmt2* mutant lines both exhibited significant reductions in plant height and culm length, and panicle length. The stacked *oscald5h1 ospmt1 ospmt2* mutant lines exhibited a similar or slightly more pronounced growth retardation phenotype compared to the *oscald5h1* and *ospmt1 ospmt2* mutant lines (**Fig. 2c; Table S3**).

### *oscald5h1 ospmt1 ospmt2* rice produces lignin lacking S units and dominated by G units

To investigate the impact of the stacked mutations in the *CAld5H* and *PMT* genes on cell wall structure, extractive-free CWR samples were prepared from mature culm tissues of the *oscald5h1*, *ospmt1 ospmt2* and *oscald5h1 ospmt1 ospmt2* mutant lines, as well as the WT control plants, and subjected to cell wall structural analyses using chemical and NMR methods.

#### Lignocellulose compositional analyses based on chemical analyses

To investigate changes in lignocellulose composition in the rice mutants, we first examined the contents of lignin, polysaccharides, and cell-wall-bound hydroxycinnamates (*p*CA and FA), in the rice culm cell walls through a series of chemical analyses. Lignin content analysis, based on the

Klason lignin assay, detected no significant changes in lignin accumulation levels in the culm cell walls of most examined mutant lines, except for one of the *oscald5h1* mutant lines, *oscald5h1-1*, which exhibited a slight reduction (**Table 1**). This result suggests that disruptions of either or both the *CAld5H* and *PMT* genes have a minor effect on the total lignin accumulation levels in the cell walls. The abundance of cell wall polysaccharides was investigated by performing neutral sugar analysis according to the two-step acid-catalyzed hydrolysis method. The amount of cellulosic glucose (glucose released from trifluoracetic acid-insoluble fractions) in the culm cell walls was not statistically different among the examined mutant and WT control lines (**Table 1**). On the other hand, the amounts of hemicellulosic sugars released by trifluoroacetic acid treatment of the culm CWR samples, including glucose, xylose, and arabinose, were all significantly reduced in the *PMT*-deficient mutant lines, i.e., *ospmt1 ospmt2*, *oscald5h1-3 ospmt1 ospmt2*, and *oscald5h1-4 ospmt1 ospmt2* (**Table 1**), suggesting that disruptions of the *PMT* genes may impair hemicellulose biosynthesis. The accumulation levels of cell-wall-bound *p*CA and FA were examined by quantifying *p*CA and FA released upon mild alkaline hydrolysis of the culm CWR samples. As expected, the *PMT*-deficient *ospmt1 ospmt2*, *oscald5h1-3 ospmt1 ospmt2*, and *oscald5h1-4 ospmt1 ospmt2* lines all displayed drastic reductions in the cell-wall-bound *p*CA content to undetectable levels. As previously reported for *ospmt1 ospmt2* (Lam *et al*., 2024), these *PMT*-deficient lines also displayed concurrent reductions in the cell-wall-bound FA content in the culm cell walls (**Table 1**).

**Table 1.**
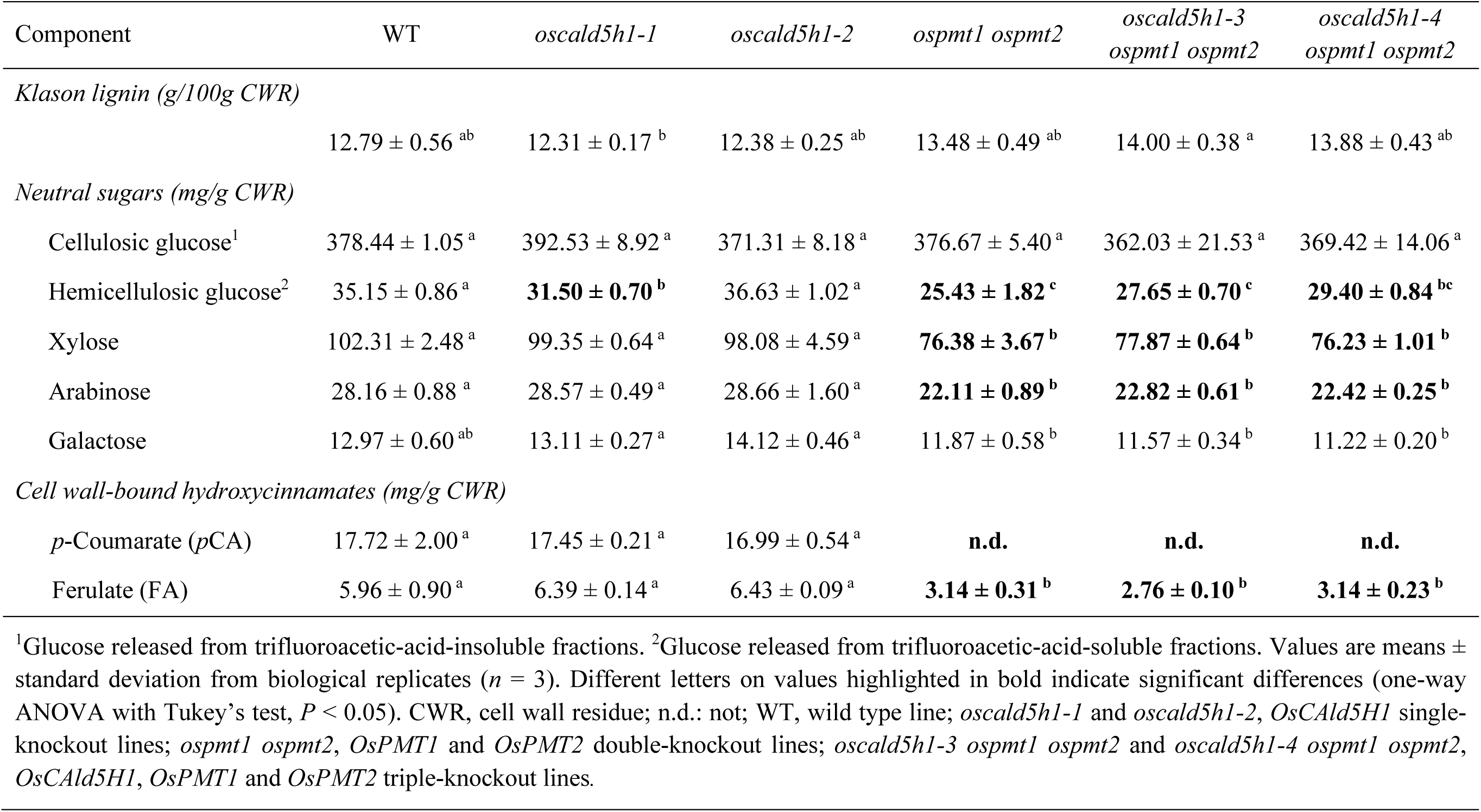
Lignocellulose composition of rice culm cell wall from wild-type, *oscald5h1*, *ospmt1 ospmt2* and *oscald5h1 ospmt1 ospmt2*.

#### Lignin compositional analyses based on DFRC and cell wall NMR

To determine the lignin monomeric composition, we first employed DFRC (Lu and Ralph, 1997; 1999; Karlen *et al*., 2018). The DFRC reactions cleave the major β–O–4-ether linkages in lignin polymers while leaving the γ-ester linkages intact, thereby releasing the γ-*p*-coumaroylated monomeric products, **G′_DFRC_** and **S′_DFRC_**, from the grass-specific γ-*p*-coumaroylated G and S units, respectively, together with non-γ-acylated monomeric products, **H_DFRC_**, **G_DFRC_**, and **S_DFRC_**, from the conventional non-γ-acylated H, G, and S units, respectively (**Fig. S1**). The *oscald5h1* lines displayed significant reductions in the S-type monomeric products, **S_DFRC_** and **S′_DFRC_**, leading to an increase in the proportion of the DFRC-based total G units over S units in the lignin polymer. However, as we previously reported (Takeda *et al*., 2019a), the *oscald5h1* lines still contained considerable amounts of **S_DFRC_** and **S′_DFRC_** (especially the latter; %**S_DFRC_** + %**S′_DFRC_** = 36–39% reduced from 51% in WT), resulting in only moderate increases in the proportion of G units (%**G_DFRC_** + %**G′_DFRC_** = 50–54% increased from 42% in WT) (**Fig. 3a; Table S4**). As also expected from our previous study (Lam *et al*., 2024), the *ospmt1 ospmt2* mutant displayed eliminations of the γ-*p*-coumaroylated monomeric products, **G′_DFRC_** and **S′_DFRC_**, to undetectable levels. Particularly due to the elimination of **S′_DFRC_**, the *ospmt1 ospmt2* mutant exhibited a significant increase of the proportion of **G_DFRC_** (%**G_DFRC_** = 62%) compared to the WT control (%**G_DFRC_**

**Fig. 3.**
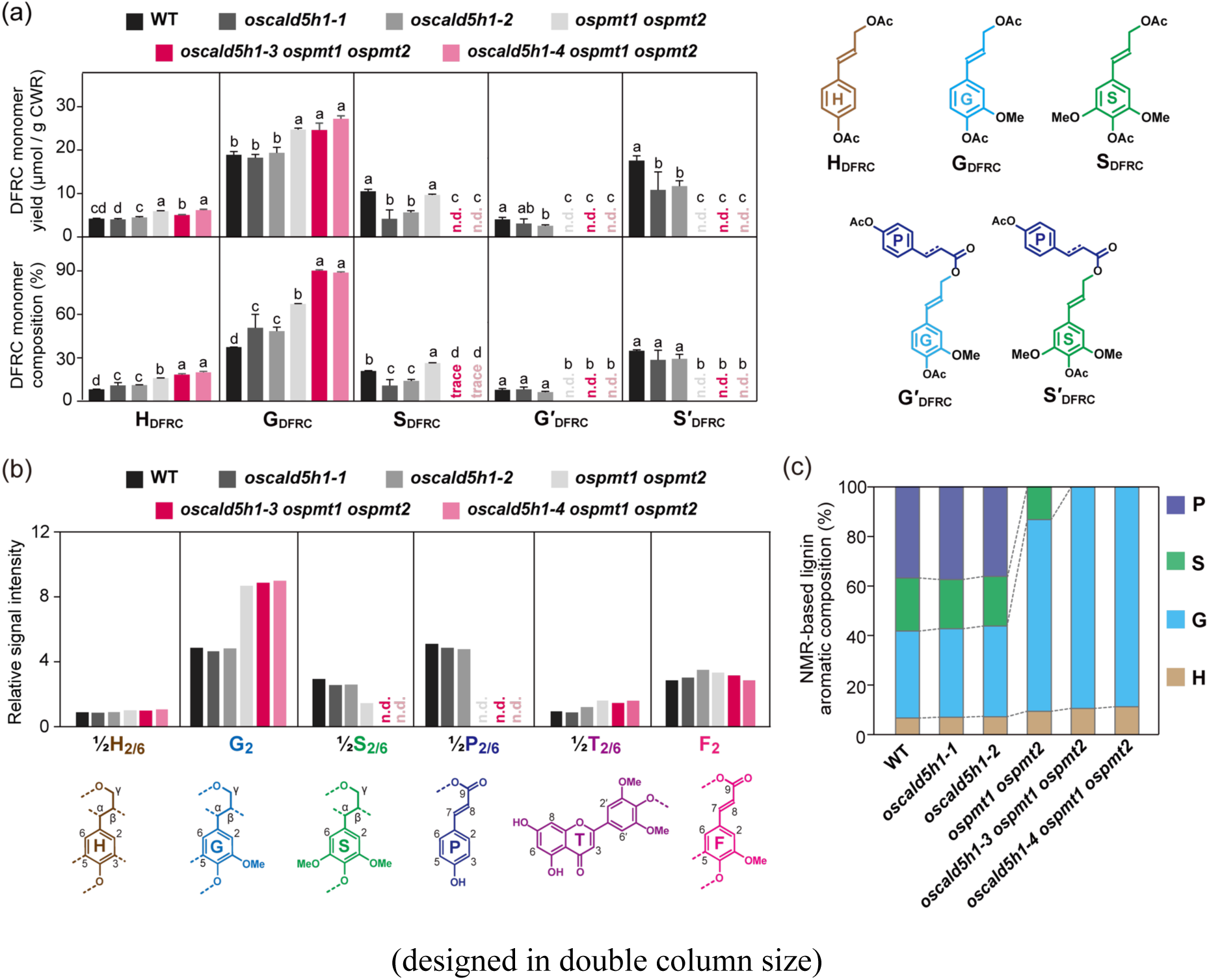
Lignin composition analysis of *CAld5H-* and *PMT*-deficient rice mutants based on DFRC and cell wall NMR. (a) Yield and composition of non-γ-acylated (**H_DFRC_**, **G_DFRC_** and **S_DFRC_**) and γ-*p*-coumaroylated (**G′_DFRC_** and **S′_DFRC_**) lignin degradation monomers released via derivatization followed by reductive cleavage (DFRC) reaction of rice culm cell wall residues (CWR). Values are means ± standard deviation from biological replicates (*n* = 3). GC-MS chromatograms and yield data of the DFRC-derived monomers are shown in **Fig. S1** and **Table S4**. (b) Normalized signal intensities of major aromatic unit signals in 2D HSQC NMR spectra of whole rice culm cell walls. The cell wall NMR spectra, complete intergradation data, and peak assignments are shown in **Fig. S2**, **Fig. S3**, and **Table S5**. (c) Estimations of the abundance of major lignin aromatic units (percentages of P, S, G, and H units relative to P + S + G + H = 100%) based on cell wall NMR integration data. n.d., not detected; S, syringyl units; G, guaiacyl units; H, *p*-hydroxyphenyl units; P, *p*-coumarate units; T, tricin units; F, ferulate units; WT, wild type line; *oscald5h1-1* and *oscald5h1-2*, *OsCAld5H1* single-knockout lines; *ospmt1 ospmt2*, *OsPMT1* and *OsPMT2* double-knockout lines; *oscald5h1-3 ospmt1 ospmt2* and *oscald5h1-4 ospmt1 ospmt2*, *OsCAld5H1*, *OsPMT1* and *OsPMT2* triple-knockout lines.

+ %**G′_DFRC_** = 42%) and *oscald5h1* mutant (%**G_DFRC_** + %**G′_DFRC_** = 50–54%) lines, but this mutant still contained a significant amount of **S_DFRC_** (%**S_DFRC_** = 24%) comparable to the WT level (%**S_DFRC_** = 19%) (**Fig. 3a; Table S4**). Strikingly, the newly developed stacked mutants, *oscald5h1-3 ospmt1 ospmt2* and *oscald5h1-4 ospmt1 ospmt2* lines, displayed eliminations of **S_DFRC_** (trace) along with **S′_DFRC_** and **G′_DFRC_** (not detected), resulting in the elimination of S-type products and a substantial increase of G-type products (%**G_DFRC_** = 81–82%) (**Fig. 3a; Table S4**). We also detected significant increases of **H_DFRC_** in the *PMT*-deficient *ospmt1 ospmt2* (%**H_DFRC_** = 15%) and *oscald5h1 ospmt1 ospmt2* (%**H_DFRC_** = 17-18%) mutant lines compared to the WT control (%**H_DFRC_** = 8%) and the *oscald5h1* mutants (%**H_DFRC_** = 10%) (**Fig. 3a; Table S4**).

The changes in the lignin aromatic composition in the rice mutants were further examined by cell wall NMR method. This technique involves acquiring 2D HSQC NMR spectra of ball-milled CWR samples using the direct dissolution/swelling method in a dimethylsulfoxide-*d*_6_/pyridine-*d*_5_ NMR solvent system (Kim and Ralph, 2010; Mansfield *et al*., 2012). The obtained HSQC spectra of the rice cell walls displayed signals from the major lignin aromatic units, including the monolignol-derived H, G, and S units (**H**, **G**, and **S**), as well as *p*CA (**P**), FA (**F**) and tricin (**T**) units, which are typical of grass lignins (**Fig. S2; Table S5**). For a semi-quantitative examination of the relative proportions of these lignin aromatic units, we performed volume integration analysis of the well-resolved lignin and *p*-hydroxycinnamate aromatic signals (**½S_2/6_**, **G_2_**, **½H_2/6_**, **½P_2/6_**, **F_2_**, and **½T_2’/6’_**) along with the major polysaccharide anomeric signals (**A_1_**, **U_1_**, **Gl_1_**, **X_1_**, **X’_1_**, and **X’’_1_**) (**Fig. S2; Table S5**). The reported signal intensity data are relative intensities normalized to the sum of the integrated signals, approximately reflecting the proportional amount of each component in the rice cell walls (**Fig. S3**).

The obtained volume integration data further corroborated the distinct lignin compositional changes observed upon the disruptions of the *CAld5H* and *PMT* genes (**Fig. 3b,c**). In agreement with the DFRC-based lignin compositional data (**Fig. 3a**), while S units exhibited only minor decreases in the *oscald5h1* mutants (**%S** relative to **P** + **S** + **G** + **H** = 100 % decreased to ∼30% from 37% in WT), they were more notably decreased in *ospmt1 ospmt2* (**%S** = 24%) and diminished to undetectable levels in *oscald5h1 ospmt1 ospmt2* along with disappearance of *p*CA units (**Fig. 3c**). Concurrently, the proportion of G units increased substantially in *ospmt1 ospmt2* (**%G** increased to 61% from 30% and 38-39% in WT and *oscald5h1*, respectively) and even more significantly in *oscald5h1 ospmt1 ospmt2* (**%G** = 82-83%). In agreement with DFRC data (**Fig. 3a**), our NMR analysis also detected proportional increases of H units in *ospmt1 ospmt2* (**%H** = 10%) and *oscald5h1 ospmt1 ospmt2* (**%H** = 10-11%) to the WT control (**%H** = 7%) and *oscald5h1* lines (**%H** = ∼7%) (**Fig. 3c**). Furthermore, tricin (**T**) signals were notably increased in all examined *CAld5H-* and *PMT*-deficient mutant lines (**Fig. 3b**). Notably, while we detected reductions of cell-wall-bound FA released by mild alkaline hydrolysis in the *PMT*-deficient mutant lines (**Table 1**), the present NMR analysis did not support these changes (**Fig. 3b**).

Collectively, the chemical and NMR data indicate that the introduction of stacked mutations in the *CAld5H* and *PMT* genes resulted in lignin predominantly composed of G units, with a virtual absence of S units. As further discussed below, these findings strongly support the existence of a CAld5H-independent parallel S lignin pathway in rice.

### In-depth structural characterizations of G-dominated lignins produced by *oscald5h1 ospmt1 ospmt2* rice

To investigate the structural features of G-dominated lignins produced by the *oscald5h1 ospmt1 ospmt2* rice mutants, dioxane/water-soluble lignin (DL) samples were prepared from the culm CWR samples of the mutants and then subjected to additional lignin structural analysis using 2D HSQC NMR and GPC. For comparison, DL samples prepared from the WT control, *oscald5h1*, and *ospmt1 ospmt2* lines were also subjected to these analyses. Additionally, to investigate the impact of the enrichment of G lignin in rice comparatively with that in Arabidopsis, DL samples were also prepared and analyzed for the *CAld5H*-deficient Arabidopsis mutant *atcald5h1* (*fah1-2*) (Meyer *et al*., 1996; 1998) and its WT control (Col-0 ecotype). Furthermore, we also included a G-dominated DL sample from a gymnosperm, pine (*P. taeda*) (Miyagawa et al., 2020), and a G-type synthetic lignin (G-DHP) prepared by *in vitro* polymerization of coniferyl alcohol (Tobimatsu *et al*., 2013).

The aromatic composition of the plant DL and G-DHP samples was established using their 2D HSQC NMR spectra (**Fig. 4a; Fig. S4a; Table S6**). Volume integration analysis of the aromatic sub-regions (δ_C_/δ_H_, 150–90/8.0–6.0 ppm) estimated the proportion of G units (the percentage of **G**_2_ relative to **½S**_2/6_ + **G**_2_ + **½H**_2/6_) to have drastically increased to ∼96% in the DL samples from the *oscald5h1 ospmt1 ospmt2* rice mutants from 45% in the WT control rice, 48– 51% in the *oscald5h1* mutant rice, and 82% in the *ospmt1 ospmt2* mutant rice. Consistent with the previous lignin composition analyses of the cell walls (**Fig. 3**), no S and *p*CA units were detected in the rice *oscald5h1 ospmt1 ospmt2* mutant lignins (**Fig. 4a; Fig. S4a**). Similarly, the proportion of G units largely increased to 99% in the *atcald5h1* Arabidopsis mutant from 82% in the WT control Arabidopsis (**Fig. 4a**). The G-dominated features of pine lignin (98%), and G-DHP (100%) were also confirmed (**Fig. 4a**).

**Fig. 4.**
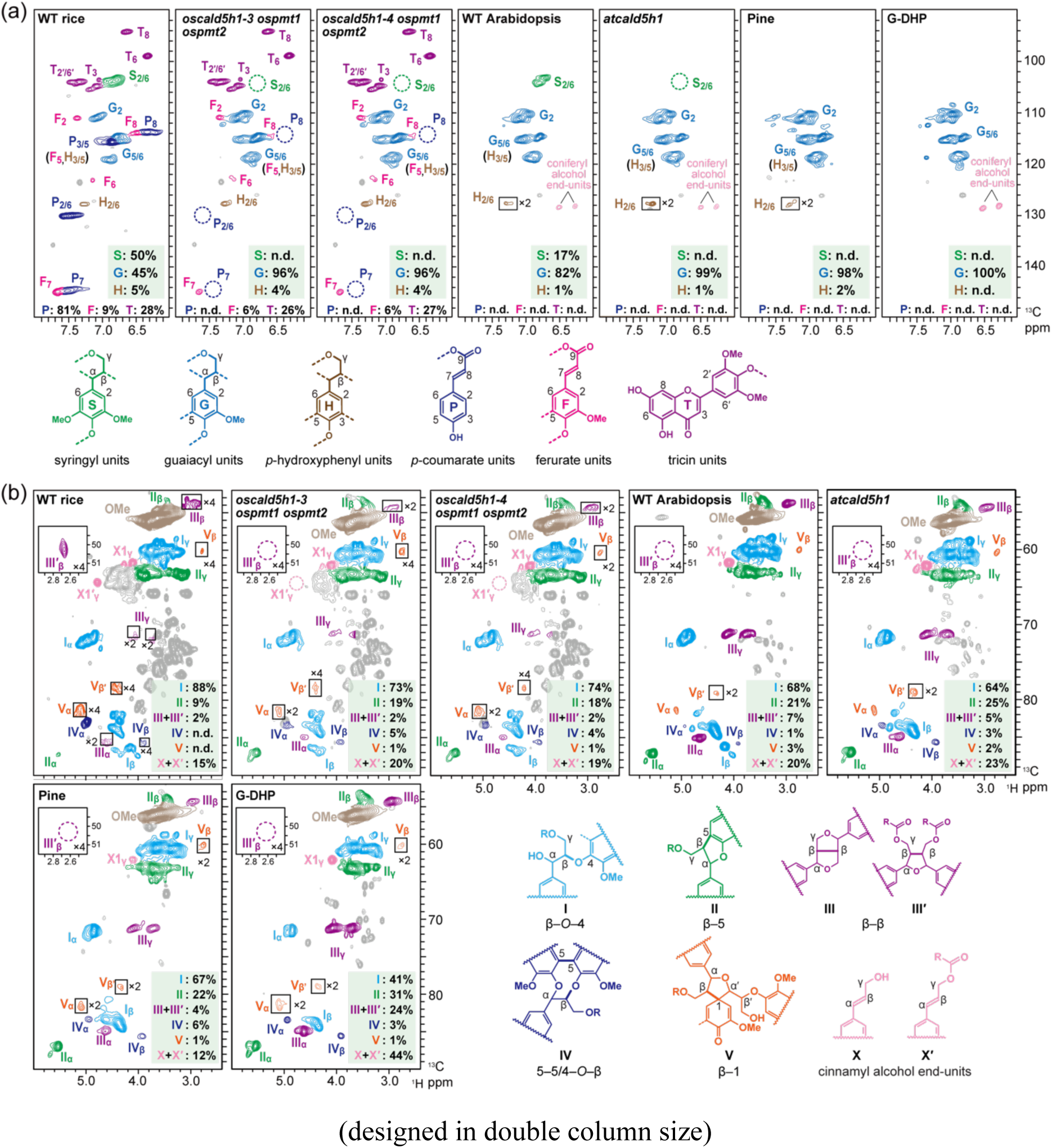
2D HSQC NMR spectra of dioxane/water-soluble lignins extracted from rice, Arabidopsis and pine, and synthetic lignin. (a) Aromatic and (b) oxygenated aliphatic sub-regions of 2D HSQC NMR spectra are displayed. The dioxane/water-soluble lignin (DL) samples were extracted from culm or stem cell walls of wild-type (WT) and *oscald5h1 ospmt1 ospmt2* mutant rice, WT (Col-0 ecotype) and *atcald5h1* mutant Arabidopsis, and pine (*Pinus taeda*). The synthetic lignin (G-DHP) was obtained by *in vitro* polymerization of coniferyl alcohol. Contour coloration matches the substructures shown in each panel. The 2D HSQC NMR spectra of DL samples from other rice lines are shown in **Fig. S4**. Peak assignments are listed in **Table S6**.Volume integrals are given for the major lignin aromatic units as percentages relative to the total of syringyl (**S**), guaiacyl (**G**), *p*-hydroxyphenyl (**H**) aromatic units (½**S_2/6_** + **G_2_** + ½**H_2/6_** = 100%) in (a) and for the major lignin inter-monomeric linkage and end-unit types as percentages relative to the total of the inter-monomeric linkage types (**I_ɑ_** + **II_ɑ_** + ½**III_ɑ_** + ½**IIIʹ_β_** + **IV_ɑ_** + **V_ɑ_** = 100%) in (b).

The oxygenated aliphatic sub-regions (δ_C_/δ_H_, 90–52/6.0–2.5 ppm) of the 2D HSQC NMR spectra displayed signals attributed to the major lignin inter-unit linkages, such as β–O–4 (**I**), β–5 (**II**), β–β (**III** and **III’**), 5–5/β–O–4 (**IV**), and β–1 (**V**), as well as those attributed to the cinnamyl alcohol end-units (**X1** and **X1’**) (**Fig. 4b; Fig. S4b; Table S6**). Due to the loss-of-function of the *PMT* genes, the signals from the tetrahydrofuran-type β–β (**IIIʹ**) units and γ-acylated cinnamyl alcohol end-units (**Xʹ**) originating from the incorporation of γ-*p*-coumaroylated monolignols (Lu and Ralph, 2002; Lam *et al*., 2024) were depleted to undetectable levels in the spectra of the DL samples from the *ospmt1 ospmt2* and *oscald5h1 ospmt1 ospmt2* rice mutants, whereas these signals were clearly seen in the spectra of the DL samples from the WT rice and the *oscald5h1* rice mutants (**Fig. 4b; Fig. S4b**). The volume integration data of the major lignin inter-unit linkage and end-unit signals (percentages relative to **I_ɑ_** + **II_ɑ_** + ½**III_ɑ_** + ½**III′_β_** + **IV_ɑ_** + **V_ɑ_** = 100%) were used to estimate the differences in the lignin linkage patterns between the lignin samples, as further reported below.

#### Impact of G lignin enrichment in rice and Arabidopsis

Compared to their corresponding WT controls, G-dominated DL samples prepared from the rice *oscald5h1 ospmt1 ospmt2* and the Arabidopsis *atcald5h1* mutants both exhibited notably reduced proportions of β–O–4 linkages (**I**) and increased proportions of β–5 (**II**) and 5–5/β–O–4 (**IV**) linkages (**Fig. 5a**). Such changes in lignin linkage distributions are typical consequences of G unit enrichment over S units in lignin (Marita *et al*., 1999; Anderson *et al*., 2015; Takeda *et al*., 2017, 2019a, 2019b). Indeed, these S/G composition-dependent lignin linkage shifts were clearly observed when comparing the linkage distributions between the DL samples from *oscald5h1*, *ospmt1 ospmt2*, and *oscald5h1 ospmt1 ospmt2* mutants (**Fig. S5**). In contrast, the rice *oscald5h1 ospmt1 ospmt2* and the Arabidopsis *atcald5h1* mutant DL samples exhibited different changes in the proportions of other linkage types compared to their corresponding WT control lignins: β–β linkages (**III** and **III’**) showed no major changes in rice but decreased in Arabidopsis; β–1 linkages (**V**) slightly increased in rice but decreased slightly in Arabidopsis; and cinnamyl alcohol end-units (**X1** and **X1’**) decreased in rice but increased in Arabidopsis (**Fig. 5a**). Consequently, the rice mutant DL samples has notably higher amounts of β–O–4 and 5–5/β–O–4 linkages and lower amounts of β–5, β–β and β–1 linkages as well as cinnamyl alcohol end-units compared to the Arabidopsis mutant DL sample (**Fig. 5a**). GPC analysis determined that the DL samples prepared from the rice *oscald5h1 ospmt1 ospmt2* and the Arabidopsis *atcald5h1* mutants both have overall similar weight-averaged molecular weight (*M*_w_) values compared to those of the corresponding WT control DL samples (**Fig. 6a,b; Table S7**), suggesting that enrichment of G units over S units has a minimal impact on the molecular weights of isolated lignins in both rice and Arabidopsis. Notably, however, the rice DL samples have considerably lower molecular weights (*M*_w_ = 8,500– 9,100 Da) compared to those of the Arabidopsis DL samples (*M*_w_ = 14,500–14,900 Da) (**Fig. 6a,b; Table S7**). Overall, these data reveal that, despite nearly identical high proportions of G units (>96%), the rice *oscald5h1 ospmt1 ospmt2* and the Arabidopsis *atcald5h1* mutant lignins exhibit notably different linkage patterns and molecular weight distributions.

**Fig. 5.**
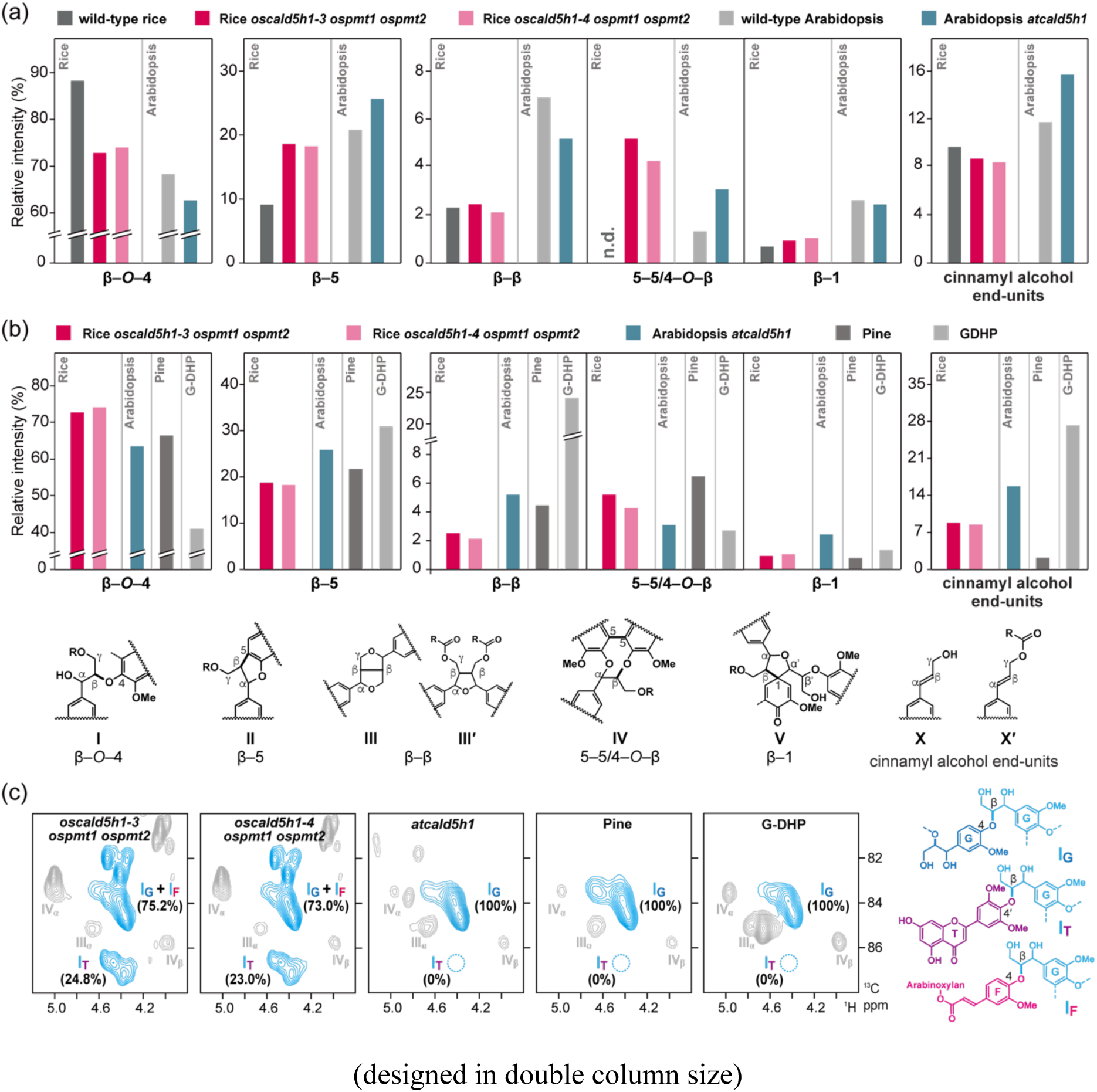
NMR-based lignin linkage distribution analysis of dioxane/water-soluble lignins extracted from rice, Arabidopsis and pine, and synthetic lignin. (a and b) Comparisons of volume integration data of lignin linkage types between dioxane/water-soluble lignin (DL) samples extracted from wild-type and G-lignin-dominated mutant lines of rice (*oscald5h1-3 ospmt1 ospmt2* and *oscald5h1-4 ospmt1 ospmt2*) and Arabidopsis (*atcald5h1*) (a) and between DL samples extracted from G-lignin-dominated rice (*oscald5h1-3 ospmt1 ospmt2* and *oscald5h1-4 ospmt1 ospmt2*) and Arabidopsis (*atcald5h1*) mutant lines, wild-type pine (*Pinus taeda*), and synthetic G lignin (G-DHP) prepared by *in vitro* polymerization of coniferyl alcohol (b). The HSQC NMR spectra of the DL samples and peak assignments are shown in **Fig. S4** and **Table S6**. Volume integral analysis was performed for the major lignin inter-monomeric linkage and end-unit types as percentages relative to the total of the analyzed inter-monomeric linkage types (**I_ɑ_** + **II_ɑ_** + ½**III_ɑ_** + **IV_ɑ_** + **V_ɑ_** = 100%). (c) Expanded β–O–4 correlation sub-regions of 2D HSQC spectra of lignin samples, highlighting **I_β_**-correlations from β–O–4 linkages linked to G (**I_G_**), tricin (**I_T_**), and ferulate (**I_F_**) units (Lan *et al*., 2018).

**Fig. 6.**
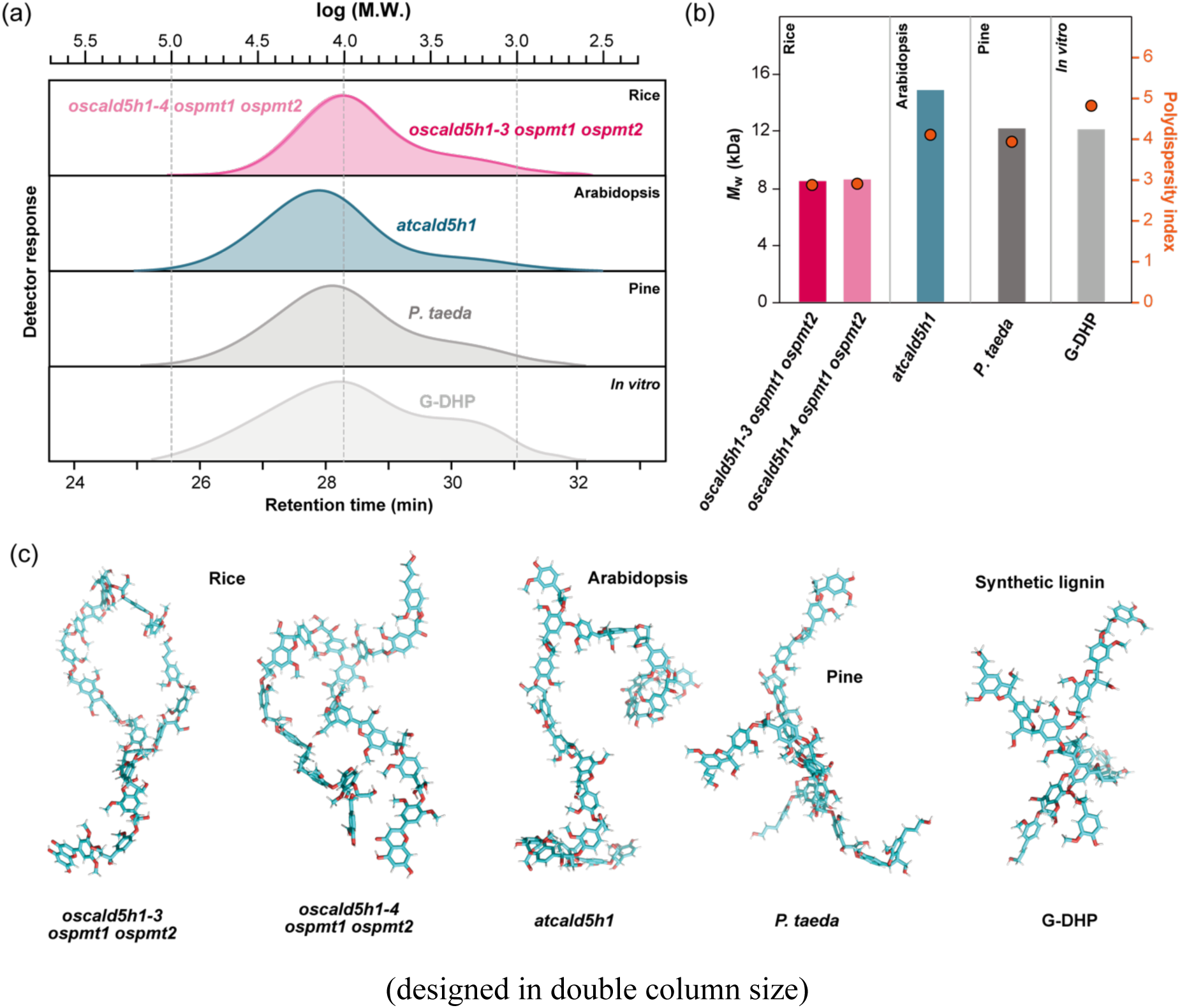
Molecular weight distributions and 3D models of dioxane/water-soluble lignins extracted from rice, Arabidopsis and pine, and synthetic lignin. Molecular weight distribution curves (a), molecular weight distribution data (b), and representative 3D structure models (c) of dioxane/water-soluble lignin (DL) samples extracted from G-lignin-dominated rice (*oscald5h1-3 ospmt1 ospmt2* and *oscald5h1-4 ospmt1 ospmt2*) and Arabidopsis (*atcald5h1*) mutant lines, wild-type pine (*Pinus taeda*), and synthetic G lignin (G-DHP) prepared by *in vitro* polymerization of coniferyl alcohol. Molecular weight distribution data were obtained by gel permeation chromatography (GPC) using polystyrene molecular weight standards and complete data are listed in **Table S7**. *M*_n_, number-averaged molecular weight; *M*_w_, weight-averaged molecular weight; PDI, polydiversity index (*M*_w_/*M*_n_). To generate 3D model structures in (c), SMILES strings were created using LigninKMC (Orella et al., 2019) based on aromatic composition and bonding patterns derived from 2D HSQC NMR and molecular weight distribution data from GPC. These SMILES strings were then used for structural modeling and optimization with LigninBuilder (Vermaas et al., 2018) and VMD (Humphrey et al., 1996) to obtain average lignin models. Finally, PyMOL (2.5 Schrödinger-LLC) was employed for visualization.

#### Comparison among G-dominated lignins produced in different plant lineages and G-DHP

The notable structural differences between the G-dominated rice and Arabidopsis mutant DL samples prompted us to expand the comparative analysis to include natural G-dominated pine DL and G-type synthetic lignin G-DHP. As previously established in earlier DHP studies (Ralph *et al*., 2004; 2009), G-DHP exhibited a markedly distinct linkage pattern compared to that of the other plant DL samples examined. It displayed a much lower content of β–O–4 linkages (**I**) and considerably higher contents of β–5 (**II**) and β–β (**III**) linkages, as well as cinnamyl alcohol end-units (**X1**), compared to those observed in the rice, Arabidopsis and pine DL samples (**Fig. 5b**). These differences can be attributed to the critical difference between the lignin polymerization environments *in vitro* and *in vivo* (Ralph *et al*., 2004; 2009), as further discussed below. Meanwhile, the G-dominated DL samples from rice, Arabidopsis and pine also exhibited notable variations in their linkage patterns. Rice DL is characterized by their higher contents of β–O–4 linkages (**I**); the Arabidopsis DL is distinguished by its higher contents of β–5 (**II**), β–β (**III**) and β–1 (**V**) linkages as well as cinnamyl alcohol end-units (**X1**); and pine DL is characterized by its higher content of 5–5/β–O–4 linkages (**IV**) compared to others (**Fig. 5b**). In addition, GPC analysis determined that the rice mutant DL samples showed notably lower molecular weights (*M*_w_ = 8,500 Da) compared to those of Arabidopsis (*M*_w_ = 14,900 Da) and pine (*M*_w_ = 12,300 Da) DL samples as well as G-DHP (*M*_w_ = 12,200) (**Fig. 6a,b; Table S7**). These structural differences between the G-dominated plant lignins (3D models illustrated in **Fig. 6c**) may reflect variations in the cell wall environment in different plant lineages that influence the polymerization of the G-type monolignol, coniferyl alcohol, during cell wall lignification, as further discussed below.

## Discussion

### Complete elimination of S lignin by stacked mutations to *CAld5H* and *PMT* provides further evidence for parallel monolignol pathways in grasses

In the present study, we demonstrated that CAld5H and PMT cooperatively act to generate S lignin in grass rice. As previously demonstrated (Takeda *et al*., 2019a), loss-of-function mutations in *OsCAld5H1*, a primary gene encoding CAld5H in rice, significantly reduce but do not eliminate S units, resulting in the persistence of substantial amounts of both non-acylated and γ-*p*-coumaroylated S units, especially the latter (**Fig. 3**). This sharply contrasts with the complete absence of S lignin in the *CAld5H*-deficient eudicot Arabidopsis mutant *atcald5h1* (Meyer *et al*., 1998; Marita *et al*., 1999; Anderson *et al*., 2015) (**Fig. 4**). Loss-of-function mutations in *OsPMT1* and *OsPMT2*, two primary genes encoding PMT in rice (Lam *et al*., 2024), also significantly reduce but do not eliminate S lignin, leaving non-acylated S units present in the mutant lignin (**Fig. 3**). Strikingly, the introduction of stacked mutations to both *CAld5H* and *PMT* genes resulted in the near-complete absence of S units in rice lignin (**Fig. 3**). This finding strongly supports the notion that rice possesses a parallel *CAld5H*-independent pathway, in which PMT plays an essential role in generating the S-type monolignol *p*-coumarate conjugate (sinapyl *p*-coumarate) alongside the conventional CAld5H-dependent pathway toward the conventional (non-γ-acylated) S-type monolignol (sinapyl alcohol) (Takeda *et al*., 2019a) (**Fig. 1**). Furthermore, this study demonstrates that targeting both *CAld5H* and *PMT* genes simultaneously provides a viable strategy for precisely manipulating lignin aromatic composition in rice and potentially other grasses.

The question of whether such CAld5H-independent monolignol pathway proposed in rice are widely present is other grass species is yet to be determined. This is particularly because most functional characterizations of *CAld5H* genes in other grasses to date have relied on incomplete downregulation of the genes using RNAi approaches. In sugarcane, RNAi-mediated downregulation of *CAld5H* genes resulted in a partial decrease in the S:G lignin unit ratio, from 61:39 to 48:52, with no detectable influence on cell-wall-bound *p*CA as determined by 2D HSQC NMR (Bewg *et al*., 2016). Similarly, in switchgrass (*Panicum virgatum L.*), RNAi-mediated downregulation of *CAld5H* genes, in combination with genes encoding 5-HYDROXYCONIFERALDEHYDE *O*-METHYLTRANSFERASE (CAldOMT), resulted in only partial reductions in S units in lignin compared to wild-type plants (Wu *et al*., 2018). In another recent study on barley (*Hordeum vulgare*), downregulation of *HvF5H1*, a primary barley *CAld5H* gene, by RNAi led to a substantial reduction in S units in lignin, with the S/G ratio reduced by up to ∼85% from 1.62 in a control plant to 0.24. Intriguingly, this reduction in S lignin content accompanied with significant reductions in cell-wall-bound *p*CA, raising questions about the existence of the parallel monolignol pathway in barley (Shafiei *et al*., 2023). However, to conclusively determine whether the parallel monolignol pathway analogous to that proposed in rice exist in these grass species, further lignin characterization of mutants with loss-of-function in *CAld5H* is necessary.

The findings in rice mutants lacking CAld5H resemble those in rice transgenic lines in which a single functional gene (*OsC3’H1*) encoding *p*-COUMAROYL ESTER 3-HYDROXYLASE (C3′H) was downregulated (Takeda *et al*., 2018; Takeda-Kimura *et al*., 2025). As C3′H catalyzes the first aromatic *m*-hydroxylation step in monolignol biosynthesis (**Fig. 1**), the downregulation of *C3′H* genes typically leads to an overaccumulation of H units with a corresponding decrease in G and S units in lignin (Franke *et al*., 2002; Ralph *et al*., 2006; Bonawitz *et al*., 2014; Takeda *et al*., 2018; Takeda-Kimura *et al*., 2025). However, similar to the increase of S units in the *CAld5H*-deficient rice (Takeda *et al*., 2019a; this study) (**Fig. 3**), the increase in H units in the *C3’H*-deficient rice only partially occurred primarily at the expense of non-γ-*p*-coumaroylated G and S units, while the levels of γ-*p*-coumaroylated G and S units remained largely unaffected (Takeda *et al*., 2018; Takeda-Kimura *et al*., 2025). This finding strongly suggests the existence of a C3′H-independent parallel pathway that synthesizes γ-*p*-coumaroylated monolignols separately from non-γ-*p*-coumaroylated monolignols, similar to the CAld5H-independent pathway explored in this study. Combined with the results of the rice mutants lacking CAld5H and PMT (Takeda *et al*., 2019a; Lam *et al*., 2024; this study), it is probable that, as illustrated in **Fig. 1**, rice may possess the capability to perform aromatic methoxylation (*m*-hydroxylation followed by *O*-methylation) of γ-*p*-coumaroylated monolignols, utilizing unidentified hydroxylase(s) [and *O*-methyltranferase(s)] and that function independently of CAld5H and C3’H. The possible role of the H-type γ-*p*-coumaroylated monolignol as a precursor for the G- and S-type γ-*p*-coumaroylated monolignols was also suggested based on the high catalytic activity of rice PMT (OsPMT1/OsAT4) toward the H-type monolignol (*p*-coumaryl alcohol) *in vitro* (Withers *et al*., 2012). However, future studies aimed at identifying missing hydroxylase and *O*-methyltransferase enzymes involved in the conversions of γ-*p*-coumaroylated monolignols are crucial for validating this possibility.

Several other studies have also implicated the possible existence of the alternative monolignol pathways towards grass-specific lignin monomers. For instance, it was reported that the function of the grass-specific bifunctional PHENYLALANINE/TYROSINE AMMONIA-LYASE (PTAL) (**Fig. 1**) in *Brachypodium distachyon* may be more specifically linked to the productions of S and *p*CA lignin units derived from sinapyl *p*-coumarate (Barros *et al*., 2016; Barros and Dixon, 2020). Two different isoforms of the rice 4-COUMAROYL-COENZYME A LIGASE (Os4CL3 and Os4CL4) (**Fig. 1**) have been reported to differentially contribute to the biosynthesis of monolignol and monolignol *p*-coumarate conjugates as well as tricin (Afifi *et al*., 2022). Moreover, OsMYB108, a rice R2R3-MYB transcription factor that negatively regulates lignin biosynthesis, is more strongly involved in the biosynthesis of the grass-specific lignin monomers, monolignol *p*-coumarate conjugates and tricin, compared to its involvement in the biosynthesis of the conventional non-γ-*p*-coumaroylated monolignols (Miyamoto *et al*., 2019; Miyamaoto *et al*., 2020).

Overall, the accumulated data collectively underscore the complexity of the metabolic network responsible for the biosynthesis of diverse lignin monomers in grasses, which substantially differs from that observed in eudicots (Barros and Dixon, 2020; Chandrakanth *et al*., 2023; Peracchi *et al*., 2024; Umezawa *et al*., 2020, 2024). A comprehensive understanding of this network requires further study of monolignol biosynthetic enzymes and genes, particularly those potentially involved in producing monolignol derivatives independent of CAld5H and C3’H in grasses (**Fig. 1**).

### Distinct structural features of G-dominated lignins from rice, Arabidopsis and pine highlight lineage-dependent lignin polymerization patterns

The successful elimination of S units along with *p*CA units in rice lignin allowed us to examine the structural characteristics of G-dominated lignins produced in rice (a herbaceous monocot grass), along with those produced in Arabidopsis (a herbaceous eudicot) and pine (a woody gymnosperm). Previous studies on lignin polymerization, primarily utilizing *in vitro* systems to prepare DHPs, have identified numerous factors that can potentially influence lignin polymer formation *in planta*. These factors include monomer supply rate, pH, ionic strength, the capacity of polymerization enzymes (laccases and peroxidases), the influence of polysaccharide matrices, etc. in lignifying cell walls where lignin monomers undergo combinatorial radical coupling (as intensely discussed, for example, in Terashima *et al*., 1995; Brunow *et al*., 1998; Cathala *et al*., 1998; Syrjänen and Brunow, 2000; Ralph *et al*., 2004; Méchin *et al*., 2007; Ralph *et al*., 2009; Tobimatsu *et al*., 2010; Hwang *et al*., 2015; Li *et al*., 2015; Tobimatsu and Schuetz, 2019; Tokunaga and Watanabe, 2023). Thus, the observed structural variations in the G-dominated plant lignins could be attributed to differences in cell wall environmental factors that modulate the polymerization of the G-type monolignol, coniferyl alcohol.

Consistent with earlier DHP studies (Sarkanen and Ludwig, 1971; Ralph *et al*., 2004; 2009), G-DHP displayed considerably different structural features compared to the G-dominated plant lignins, with a markedly low level of β–O–4 linkages and high levels of β–5, β–β, and cinnamyl alcohol end-unit linkages (**Fig. 5**). These distinct differences between DHP and plant lignins can be primarily attributed to the over-representation of dehydrodimerization reactions over cross-coupling reactions in typical DHP preparation conditions. While dehydrodimerization reactions of coniferyl alcohol typically yield β–O–4, β–5 and β–β dimers at comparable levels, cross-coupling reactions of coniferyl alcohol with growing lignin polymers favor the formation of β–O–4 linkages (Syrjänen and Brunow, 2000; Ralph *et al*., 2004; 2009). Nevertheless, despite exhibiting general structural similarities with each other when compared to G-DHP, our NMR and GPC analyses detected notable structural differences among the three G-dominated plant lignins in terms of their lignin linkage distributions (**Fig. 5**) and molecular weight distributions (**Fig. 6**).

The G-dominated lignins isolated from the rice *oscald5h1 ospmt1 ospmt2* mutants were characterized by their relatively higher contents of β–O–4 linkages and also their lower molecular weights compared to those from Arabidopsis and pine (**Fig. 5**; **Fig. 6**). These differences may be attributed, at least in part, to the presence of grass-specific tricin and ferulate nucleation sites within grass cell walls, potentially influencing lignin polymerization initiation. Tricin, a canonical lignin monomer in grasses, has been shown to exclusively cross-couple with monolignols and γ-acylated monolignols via the β–O–4-type coupling mode (4ʹ–O–β-coupling) both *in vivo* and *in vitro* (del Río *et al*., 2012; 2020; Lan *et al*., 2015; Elder *et al*., 2020). Since tricin is unable to participate in homocoupling or direct cross-coupling with growing lignin polymers, it preferentially incorporates into the starting point of nascent lignin polymer chains (via cross-coupling with other lignin monomers), thereby serving as an initiation site for lignin polymerization (Lan *et al*., 2015; del Río *et al*., 2020; Berstis *et al*., 2021; Lam *et al*., 2021). Similarly, ferulate, which is predominantly esterified to arabinoxylan, serves as an initiation site for lignin polymerization and preferentially cross-couples with monolignols via the β–O–4 linkage during cell wall lignification in grasses (Bunzel *et al*., 2004; Ralph, 2010). Therefore, the abundant presence of tricin and ferulate initiation sites in grass cell walls may contribute to the increased formation of β–O–4 linkages in lignin, as observed in the G-dominated rice lignin in the present study. Supporting this idea, we found that a significant portion of the β–O–4 linkages in the rice mutant lignins originated from the cross-coupling of coniferyl alcohol with tricin (**Fig. 5c**). The abundant presence of tricin and ferulate initiation sites may also contribute to limiting the growth of lignin polymer chains in grass cell walls. This occurs because the available pool of coniferyl alcohol is distributed among a larger number of growing polymer chains originating from abundant initiation sites, reducing the amount of coniferyl alcohol available for the elongation of each individual chain.

The G-dominated lignin from the Arabidopsis *atcald5h1* mutant exhibited “bulk polymer” features more closely resembling G-DHP, with a lower content of β–O–4 linkages and higher contents of β–5, β–β, and cinnamyl alcohol end-unit linkages compared to the other G-dominated lignins from rice and pine (**Fig. 5**). Since such “bulk polymer” features are well-known to be associated with rapid monomer supply rates during lignification (Sarkanen and Ludwig, 1971; Ralph *et al*., 2004; 2009; Tokunaga and Watanabe, 2023), the relatively fast lignification process during stem development in Arabidopsis may promote the formation of bulk-polymer-type linkages in lignin. On the other hand, the G-dominated lignin from pine exhibited higher amounts of 5–5/β–O–4 linkages (**Fig. 5**). Since the formation of 5–5 linkages primarily occurs through cross-coupling reactions between polymer/oligomer chains during the later stages of lignin polymerization (Sarkanen and Ludwig, 1971; Brunow *et al*., 1998; Ralph *et al*., 2004; 2009; Tokunaga and Watanabe, 2023), this observation suggests that, in contrast to the rapid Arabidopsis lignification as proposed above, the relatively slower, long-term lignification process during pine wood formation might favor the preferential occurrence of 5–5 linkages in pine lignin.

One of the apparent differences in lignin polymerization among the three plant species examined in this study is the distinct composition of their cell wall polysaccharide matrices. Specifically, the cell wall polysaccharide matrices in rice (grass), Arabidopsis (eudicot), and pine (gymnosperm) are predominantly composed of arabinoxylan, glucuronoxylan, and glucomannan, respectively (Scheller and Ulvskov, 2010). As lignin polymerization *in planta* occurs after the deposition of polysaccharides in developing cell walls, pre-existing polysaccharide matrices may act as scaffolds for lignin polymerization, providing specific local environments wherein lignin monomers undergo radical coupling reactions. Indeed, prior research has demonstrated that DHP formation can be affected by the presence of various polysaccharides in the polymerization system (Terashima *et al*., 1995; Cathala and Monties, 2001; Nakamura *et al*., 2006; Barakat *et al*., 2007; Li *et al*., 2015; Warinowski *et al*., 2016; Aminzadeh *et al*., 2017). However, the influence of varying polysaccharide matrices on lignin linkage patterns is still not well-defined. It would therefore be intriguing to revisit DHP preparations in the presence of different polysaccharide matrices and analyze the resultant DHP structures using advanced lignin characterization techniques. Further in-depth comparative structural characterizations of lignins with well-defined monomer composition, generated both *in vivo* and *in vitro*, will be crucial for advancing our understanding of the factors controlling lignin polymerization *in planta*. Such studies will ultimately enhance our ability to manipulate lignin structure for improved lignocellulose utilization.

## Acknowledgments

We thank Prof. Hironori Kaji and Ms. Ayaka Maeno (ICR, Kyoto University) for their assistance in NMR analysis. This work was supported in part by grants from the Japan Society for the Promotion of Science (grant nos. KAKENHI #JP20H03044 and #JP24K01827). S.Y. acknowledges the JSPS fellowship program (grant no. #JP22J13457). A part of this study was conducted using the facilities at the DASH/FBAS, RISH, Kyoto University, and the NMR spectrometer at JURC, ICR, Kyoto University.

## Competing interests

The authors declare that they have no competing interests.

## Data availability

The authors confirm that the data supporting the findings of this study are available within the article and its supplementary materials.

## Author contributions

P.J., T.U. and Y.T. conceived research. P.J., O.A.A, S.Y., Y.O., K.O. and Y.T. performed experiments, and analyzed the data. P.J. and Y.T. wrote the manuscript with help from all the others.

## Supporting Information

**Fig. S1** DFRC-derived lignin monomers released from rice cell walls.

**Fig. S2** 2D HSQC NMR spectra of whole rice cell walls.

**Fig. S3** Volume integration analysis of 2D HSQC NMR spectra of rice cell walls.

**Fig. S4** 2D HSQC NMR spectra of dioxane/water-soluble lignins.

**Fig. S5** NMR-based lignin linkage distribution analysis of dioxane/water-soluble lignins.

**Table S1** Primers and oligonucleotides used in this study.

**Table S2** Accession numbers of *CAld5H* and *PMT* genes studied.

**Table S3** Growth characteristics of rice mutants.

**Table S4** Yield and composition of DFRC-derived lignin monomers.

**Table S5** Peak assignments in 2D HSQC NMR spectra of rice cell walls.

**Table S6** Peak assignments in 2D HSQC NMR spectra of dioxane/water-soluble lignins.

**Table S7** Molecular weight distribution data of dioxane/water-soluble lignins and synthetic lignin.

## Notes

### Competing Interest Statement

The authors have declared no competing interest.

## References

Adler E. 1977. Lignin chemistry-past, present and future. Wood Science and Technology 11: 169–218.

Afifi OA, Tobimatsu Y, Lam PY, Martin AF, Miyamoto T, Osakabe Y, Osakabe K, Umezawa T. 2022. Genome-edited rice deficient in two *4-COUMARATE:COENZYME A LIGASE* genes displays diverse lignin alterations. Plant Physiology 190: 2155–2172.

Aminzadeh S, Zhang LM, Henriksson G 2017. A possible explanation for the structural inhomogeneity of lignin in LCC networks. Wood Science and Technology 51: 1365–1376.

Anderson NA, Tobimatsu Y, Ciesielski PN, Ximenes E, Ralph J, Donohoe BS, Ladisch M, Chapple C. 2015. Manipulation of Guaiacyl and Syringyl Monomer Biosynthesis in an Arabidopsis Cinnamyl Alcohol Dehydrogenase Mutant Results in Atypical Lignin Biosynthesis and Modified Cell Wall Structure. The Plant Cell 27: 2195–2209.

Barakat A, Chabbert B, Cathala B. 2007. Effect of reaction media concentration on the solubility and the chemical structure of lignin model compounds. Phytochemistry 68: 2118–2125.

Barros J, Dixon RA. 2020. Plant phenylalanine/tyrosine ammonia-lyases. Trends in Plant Science 25: 66–79.

Barros J, Serrani-Yarce JC, Chen F, Baxter D, Venables BJ, Dixon RA. 2016. Role of bifunctional ammonia-lyase in grass cell wall biosynthesis. Nature plants 2: 1–9.

Berstis L, Elder T, Dixon R, Crowley M, Beckham GT. 2021. Coupling of flavonoid initiation sites with monolignols studied by density functional theory. ACS Sustainable Chemistry & Engineering 9: 1518–1528.

Bewg WP, Poovaiah C, Lan W, Ralph J, Coleman HD. 2016. RNAi downregulation of three key lignin genes in sugarcane improves glucose release without reduction in sugar production. Biotechnology for Biofuels 9: 1–13.

Bhatia R, Gallagher JA, Gomez LD, Bosch M. 2017. Genetic engineering of grass cell wall polysaccharides for biorefining. Plant Biotechnology Journal 15: 1071–1092.

Bjorkman A. 1957. Lignin and lignin-carbohydrate complexes. Industrial & Engineering Chemistry 49: 1395–1398.

Bonawitz ND, Kim JI, Tobimatsu Y, Ciesielski PN, Anderson NA, Ximenes E, Maeda J, Ralph J, Donohoe BS, Ladisch M, Chapple C. 2014. Disruption of Mediator rescues the stunted growth of a lignin-deficient *Arabidopsis* mutant. Nature 509: 376–380.

Brunow G, Kilpeläinen I, Sipilä J, Syrjänen K, Karhunen P, Setälä H, Rummakko P. 1998. Oxidative coupling of phenols and the biosynthesis of lignin. Lignin and Lignan Biosynthesis, 131–147.

Bunzel M, Ralph J, Lu F, Hatfield RD, Steinhart H. 2004. Lignins and ferulate− coniferyl alcohol cross-coupling products in cereal grains. Journal of Agricultural and Food Chemistry 52: 6496–6502.

Cathala B, Saake B, Faix O, Monties B. 1998. Evaluation of the reproducibility of the synthesis of dehydrogenation polymer models of lignin. Polymer Degradation and Stability 59: 65–69.

Cathala B, Monties B. 2001. Influence of pectins on the solubility and the molar mass distribution of dehydrogenative polymers (DHPs, lignin model compounds). International Journal of Biological Macromolecules 29: 45–51.

Chandrakanth NN, Zhang C, Freeman J, De Souza WR, Bartley LE, Mitchell RA. 2023. Modification of plant cell walls with hydroxycinnamic acids by BAHD acyltransferases. Frontiers in Plant Science 13: 1088879.

Davis R, Kataria R, Cerrone F, Woods T, Kenny S, O’donovan A, Guzik M, Shaikh H, Duane G, Gupta VK. 2013. Conversion of grass biomass into fermentable sugars and its utilization for medium chain length polyhydroxyalkanoate (mcl-PHA) production by *Pseudomonas* strains. Bioresource Technology 150: 202–209.

Del Río JC, Rencoret J, Gutiérrez A, Elder T, Kim H, Ralph J. 2020. Lignin monomers from beyond the canonical monolignol biosynthetic pathway: another brick in the wall. ACS Sustainable Chemistry & Engineering 8: 4997–5012.

Del Río JC, Rencoret J, Prinsen P, Martínez ÁT, Ralph J, GutiéRrez A. 2012. Structural characterization of wheat straw lignin as revealed by analytical pyrolysis, 2D-NMR, and reductive cleavage methods. Journal of Agricultural and Food Chemistry 60: 5922–5935.

Edmunds CW, Peralta P, Kelley SS, Chiang VL, Sharma-Shivappa RR, Davis MF, Harman-Ware AE, Sykes RW, Gjersing E, Cunningham MW. 2017. Characterization and enzymatic hydrolysis of wood from transgenic *Pinus taeda* engineered with syringyl lignin or reduced lignin content. Cellulose 24: 1901–1914.

Elder T, Del Río JC, Ralph J, Rencoret J, Kim H, Beckham GT, Crowley MF. 2020. Coupling and reactions of lignols and new lignin monomers: A density functional theory study. ACS Sustainable Chemistry & Engineering 8: 11033–11045.

Franke R, Mcmichael CM, Meyer K, Shirley AM, Cusumano JC, Chapple C. 2000. Modified lignin in tobacco and poplar plants over-expressing the Arabidopsis gene encoding ferulate 5-hydroxylase. The Plant Journal 22: 223–234.

Franke R, Hemm MR, Denault JW, Ruegger MO, Humphreys JM, Chapple C. 2002. Changes in secondary metabolism and deposition of an unusual lignin in the *ref8* mutant of *Arabidopsis*. The Plant Journal 30: 47–59.

Freudenberg K, Neish AC. 1968. Constitution and biosynthesis of lignin, Springer Berlin, Heidelberg.

Hatfield RD, Jung HJG, Ralph J, Buxton DR, Weimer PJ. 1994. A comparison of the insoluble residues produced by the Klason lignin and acid detergent lignin procedures. Journal of the Science of Food and Agriculture 65: 51–58.

Hiei Y, Ohta S, Komari T, Kumashiro T. 1994. Efficient transformation of rice (*Oryza sativa* L.) mediated by *Agrobacterium* and sequence analysis of the boundaries of the T-DNA. The Plant Journal 6: 271–282.

Higuchi T. 1985. Biosynthesis of lignin. In: Biosynthesis and Biodegradation of Wood Components. Higuchi T. eds. Academic Press. pp141–160.

Humphrey W, Dalke A, Schulten K. 1996. VMD - Visual Molecular Dynamics. Journal of Molecular Graphics 14: 33–38.

Humphreys JM, Hemm MR, Chapple C. 1999. New routes for lignin biosynthesis defined by biochemical characterization of recombinant ferulate 5-hydroxylase, a multifunctional cytochrome P450-dependent monooxygenase. Proceedings of the National Academy of Sciences 96: 10045–10050.

Hwang H, Moon S-J, Won K, Kim YH, Choi JW. 2015. Parameters affecting *in vitro* monolignol couplings during dehydrogenative polymerization in the presence of peroxidase and H_2_O_2_. Journal of Industrial and Engineering Chemistry 26: 390–395.

Karlen SD, Free HC, Padmakshan D, Smith BG, Ralph J, Harris PJ. 2018. Commelinid monocotyledon lignins are acylated by *p*-coumarate. Plant Physiology 177: 513–521.

Kim H, Ralph J. 2010. Solution-state 2D NMR of ball-milled plant cell wall gels in DMSO-*d*_6_/pyridine-*d*_5_. Organic & Biomolecular Chemistry 8: 576–591.

Koshiba T, Yamamoto N, Tobimatsu Y, Yamamura M, Suzuki S, Hattori T, Mukai M, Noda S, Shibata D, Sakamoto M. 2017. MYB-mediated upregulation of lignin biosynthesis in *Oryza sativa* towards biomass refinery. Plant Biotechnology 34: 7–15.

Lal R. 2005. World crop residues production and implications of its use as a biofuel. Environment International 31: 575–584.

Lam LPY, Tobimatsu Y, Suzuki S, Tanaka T, Yamamoto S, Takeda-Kimura Y, Osakabe Y, Osakabe K, Ralph J, Bartley LE, et al. 2024. Disruption of *p*-coumaroyl-CoA:monolignol transferases in rice drastically alters lignin composition. Plant Physiology 194: 832–848.

Lam PY, Lui AC, Wang L, Liu H, Umezawa T, Tobimatsu Y, Lo C. 2021. Tricin biosynthesis and bioengineering. Frontiers in Plant Science 12: 733198.

Lam PY, Tobimatsu Y, Takeda Y, Suzuki S, Yamamura M, Umezawa T, Lo C. 2017. Disrupting Flavone Synthase II Alters Lignin and Improves Biomass Digestibility. Plant Physiology 174: 972–985.

Lan W, Lu F, Regner M, Zhu Y, Rencoret J, Ralph SA, Zakai UI, Morreel K, Boerjan W, Ralph J. 2015. Tricin, a flavonoid monomer in monocot lignification. Plant physiology 167: 1284–1295.

Lan W, Yue F, Rencoret J, Del Río JC, Boerjan W, Lu F, Ralph J. 2018. Elucidating tricin-lignin structures: assigning correlations in HSQC spectra of monocot lignins. Polymers 10: 916.

Li L, Zhou Y, Cheng X, Sun J, Marita JM, Ralph J, Chiang VL. 2003. Combinatorial modification of multiple lignin traits in trees through multigene cotransformation. Proceedings of the National Academy of Sciences 100: 4939–4944.

Li Q, Koda K, Yoshinaga A, Takabe K, Shimomura M, Hirai Y, Tamai Y, Uraki Y. 2015. Dehydrogenative polymerization of coniferyl alcohol in artificial polysaccharides matrices: effects of xylan on the polymerization. Journal of Agricultural and Food Chemistry 63: 4613–4620.

Liu H, Ding Y, Zhou Y, Jin W, Xie K, Chen L. 2017. CRISPR-P 2.0: an improved CRISPR-Cas9 tool for genome editing in plants. Molecular plant 10: 530–532.

Lu F, Ralph J. 1997. Derivatization followed by reductive cleavage (DFRC method), a new method for lignin analysis: protocol for analysis of DFRC monomers. Journal of Agricultural and Food Chemistry 45: 2590–2592.

Lu F, Ralph J. 1999. Detection and determination of *p*-coumaroylated units in lignins. Journal of Agricultural and Food Chemistry 47: 1988–1992.

Lu F, Ralph J. 2002. Preliminary evidence for sinapyl acetate as a lignin monomer in kenaf. Chemical Communications 1: 90–91.

Mansfield SD, Kim H, Lu F, Ralph J. 2012. Whole plant cell wall characterization using solution-state 2D NMR. Nature Protocols 7: 1579–1589.

Marita JM, Ralph J, Hatfield RD, Chapple C. 1999. NMR characterization of lignins in Arabidopsis altered in the activity of ferulate 5-hydroxylase. Proceedings of the National Academy of Sciences 96: 12328–12332.

Martin AF, Tobimatsu Y, Lam PY, Matsumoto N, Tanaka T, Suzuki S, Kusumi R, Miyamoto T, Takeda-Kimura Y, Yamamura M. 2023. Lignocellulose molecular assembly and deconstruction properties of lignin-altered rice mutants. Plant Physiology 191: 70–86.

Méchin V, Baumberger S, Pollet B, Lapierre C. 2007. Peroxidase activity can dictate the *in vitro* lignin dehydrogenative polymer structure. Phytochemistry 68: 571–579.

Meyer K, Cusumano JC, Somerville C, Chapple C. 1996. Ferulate-5-hydroxylase from Arabidopsis thaliana defines a new family of cytochrome P450-dependent monooxygenases. Proceedings of the National Academy of Sciences 93: 6869–6874.

Meyer K, Shirley AM, Cusumano JC, Bell-Lelong DA, Chapple C. 1998. Lignin monomer composition is determined by the expression of a cytochrome P450-dependent monooxygenase in *Arabidopsis*. Proceedings of the National Academy of Sciences 95: 6619–6623.

Miyagawa Y, Tobimatsu Y, Lam PY, Mizukami T, Sakurai S, Kamitakahara H, Takano T. 2020. Possible mechanisms for the generation of phenyl glycoside-type lignin-carbohydrate linkages in lignification with monolignol glucosides. The Plant Journal 104: 156–170.

Miyamoto T, Takada R, Tobimatsu Y, Takeda Y, Suzuki S, Yamamura M, Osakabe K, Osakabe Y, Sakamoto M, Umezawa T. 2019. OsMYB 108 loss-of-function enriches *p*-coumaroylated and tricin lignin units in rice cell walls. The Plant Journal 98: 975–987.

Miyamoto T, Tobimatsu Y, Umezawa T. 2020. MYB-mediated regulation of lignin biosynthesis in grasses. Current Plant Biology 24: 100174.

Mullet J, Morishige D, Mccormick R, Truong S, Hilley J, Mckinley B, Anderson R, Olson SN, Rooney W. 2014. Energy Sorghum—a genetic model for the design of C4 grass bioenergy crops. Journal of Experimental Botany 65: 3479–3489.

Nakamura R, Matsushita Y, Umemoto K, Usuki A, Fukushima K. 2006. Enzymatic polymerization of coniferyl alcohol in the presence of cyclodextrins. Biomacromolecules 7: 1929–1934.

Orella MJ, Gani TZ, Vermaas JV, Stone ML, Anderson EM, Beckham GT, Brushett FR, Román-Leshkov Y. 2019. Lignin-KMC: A toolkit for simulating lignin biosynthesis. ACS Sustainable Chemistry & Engineering 7: 18313–18322.

Osakabe K, Tsao CC, Li L, Popko JL, Umezawa T, Carraway DT, Smeltzer RH, Joshi CP, Chiang VL. 1999. Coniferyl aldehyde 5-hydroxylation and methylation direct syringyl lignin biosynthesis in angiosperms. Proceedings of the National Academy of Sciences 96: 8955–8960.

Peracchi LM, Panahabadi R, Barros-Rios J, Bartley LE, Sanguinet KA. 2024. Grass lignin: biosynthesis, biological roles, and industrial applications. Frontiers in Plant Science 15: 1343097.

Ralph J, Lundquist K, Brunow G, Lu F, Kim H, Schatz PF, Marita JM, Hatfield RD, Ralph SA, Christensen JH, et al. 2004. Lignins: natural polymers from oxidative coupling of 4-hydroxyphenyl-propanoids. Phytochemistry Reviews 3: 29–60.

Ralph J, Akiyama T, Kim H, Lu F, Schatz PF, Marita JM, Ralph SA, Reddy MSS, Chen F, Dixon RA. 2006. Effects of coumarate 3-hydroxylase down-regulation on lignin structure. Journal of Biological Chemistry 281: 8843–8853.

Ralph J, Brunow G, Harris PJ, Dixon RA, Schatz PF, Boerjan W. 2009. Lignification: are lignins biosynthesized via simple combinatorial chemistry or via proteinaceous control and template replication. Recent Advances in Polyphenol Research 1: 36–66.

Ralph J. 2010. Hydroxycinnamates in lignification. Phytochemistry Reviews 9: 65–83.

Ralph J, Lapierre C, Boerjan W. 2019. Lignin structure and its engineering. Current Opinion in Biotechnology 56: 240–249.

Ralph SA, Ralph J, Lu F. 2024. NMR database of lignin and cell wall model compounds. Available at 10.11578/2409191.

Reddy MS, Chen F, Shadle G, Jackson L, Aljoe H, Dixon RA. 2005. Targeted down-regulation of cytochrome P450 enzymes for forage quality improvement in alfalfa (*Medicago sativa* L.). Proceedings of the National Academy of Sciences 102: 16573–16578.

Sarkanen KV, Ludwig CH. 1971. Lignins: occurrence, formation, structure and reactions. Wiley Interscience, New York.

Scheller HV, Ulvskov P. 2010. Hemicelluloses. Annual review of plant biology 61: 263–289.

Shafiei R, Hooper M, Mcclellan C, Oakey H, Stephens J, Lapierre C, Tsuji Y, Goeminne G, Vanholme R, Boerjan W, et al. 2023. Downregulation of barley ferulate 5-hydroxylase dramatically alters straw lignin structure without impact on mechanical properties. Frontiers in Plant Science 13: 1125003.

Stewart JJ, Akiyama T, Chapple C, Ralph J, Mansfield SD. 2009. The effects on lignin structure of overexpression of ferulate 5-hydroxylase in hybrid poplar^1^. Plant physiology 150: 621–635.

Syrjänen K, Brunow G. 2000. Regioselectivity in lignin biosynthesis. The influence of dimerization and cross-coupling. Journal of the Chemical Society, Perkin Transactions 1: 183–187.

Takeda Y, Koshiba T, Tobimatsu Y, Suzuki S, Murakami S, Yamamura M, Rahman MM, Takano T, Hattori T, Sakamoto M, et al. 2017. Regulation of *CONIFERALDEHYDE 5-HYDROXYLASE* expression to modulate cell wall lignin structure in rice. Planta 246: 337–349.

Takeda Y, Tobimatsu Y, Karlen SD, Koshiba T, Suzuki S, Yamamura M, Murakami S, Mukai M, Hattori T, Osakabe K, et al. 2018. Downregulation of *p-COUMAROYL ESTER 3-HYDROXYLASE* in rice leads to altered cell wall structures and improves biomass saccharification. The Plant Journal 95: 796–811.

Takeda Y, Suzuki S, Tobimatsu Y, Osakabe K, Osakabe Y, Ragamustari SK, Sakamoto M, Umezawa T. 2019a. Lignin characterization of rice *CONIFERALDEHYDE 5-HYDROXYLASE* loss-of-function mutants generated with the CRISPR/Cas9 system. The Plant Journal 97: 543–554.

Takeda Y, Tobimatsu Y, Yamamura M, Takano T, Sakamoto M, Umezawa T. 2019b. Comparative evaluations of lignocellulose reactivity and usability in transgenic rice plants with altered lignin composition. Journal of Wood Science 65: 1–11.

Takeda-Kimura Y, Mori T, Suzuki S, Sakamoto M, Saito K, Nakabayashi R, Tobimatsu Y, Umezawa T. 2025. Altered development and lignin deposition in rice p-*COUMAROYL ESTER 3-HYDROXYLASE* loss-of-function mutants. The Plant Journal 121: e70039.

Terashima N, Atalla R, Ralph S, Landucci LL, Lapierre C, Monties B. 1995. New preparations of lignin polymer models under conditions that approximate cell wall lignification. I. Synthesis of novel lignin polymer models and their structural characterization by ^13^C NMR. Holzforschung 49: 521–527.

Tetreault HM, Gries T, Palmer NA, Funnell-Harris DL, Sato S, Ge Z, Sarath G, Sattler SE. 2020. Overexpression of ferulate 5-hydroxylase increases syringyl units in *Sorghum bicolor*. Plant molecular biology 103: 269–285.

Tobimatsu Y, Chen F, Nakashima J, Escamilla-Treviño LL, Jackson L, Dixon RA, Ralph J. 2013. Coexistence but independent biosynthesis of catechyl and guaiacyl/syringyl lignin polymers in seed coats. The Plant Cell 25: 2587–2600.

Tobimatsu Y, Takano T, Kamitakahara H, Nakatsubo F. 2010. Reactivity of syringyl quinone methide intermediates in dehydrogenative polymerization. Part 2: pH effect in horseradish peroxidase-catalyzed polymerization of sinapyl alcohol. Holzforschung 64: 183–192.

Tobimatsu Y, Schuetz M. 2019. Lignin polymerization: how do plants manage the chemistry so well? Current Opinion in Biotechnology 56: 75–81.

Toda E, Koiso N, Takebayashi A, Ichikawa M, Kiba T, Osakabe K, Osakabe Y, Sakakibara H, Kato N, Okamoto T. 2019. An efficient DNA-and selectable-marker-free genome-editing system using zygotes in rice. Nature Plants 5: 363–368.

Tokunaga Y, Watanabe T. 2023. An investigation of the factors controlling the chemical structures of lignin dehydrogenation polymers. Holzforschung 77: 51–62.

Tye YY, Lee KT, Abdullah WNW, Leh CP. 2016. The world availability of non-wood lignocellulosic biomass for the production of cellulosic ethanol and potential pretreatments for the enhancement of enzymatic saccharification. Renewable and Sustainable Energy Reviews 60: 155–172.

Umezawa T. 2024. Metabolic engineering of *Oryza sativa* for lignin augmentation and structural simplification. Plant Biotechnology 41: 89–101.

Umezawa T, Tobimatsu Y, Yamamura M, Miyamoto T, Takeda Y, Koshiba T, Takada R, Lam PY, Suzuki S, Sakamoto M. 2020. Lignin metabolic engineering in grasses for primary lignin valorization. Lignin 1: 30–41.

Vermaas JV, Dellon LD, Broadbelt LJ, Beckham GT, Crowley MF. 2018. Automated transformation of lignin topologies into atomic structures with LigninBuilder. ACS Sustainable Chemistry & Engineering 7: 3443–3453.

Wagner A, Tobimatsu Y, Phillips L, Flint H, Geddes B, Lu F, Ralph J. 2015. Syringyl lignin production in conifers: Proof of concept in a Pine tracheary element system. Proceedings of the National Academy of Sciences 112: 6218–6223.

Warinowski T, Koutaniemi S, Karkonen A, Sundberg I, Toikka M, Simola LK, Kilpelainen I, Teeri TH. 2016. Peroxidases bound to the growing lignin polymer produce natural like extracellular lignin in a cell culture of Norway spruce. Frontiers in Plant Science 7: 1523.

Withers S, Lu F, Kim H, Zhu Y, Ralph J, Wilkerson CG. 2012. Identification of grass-specific enzyme that acylates monolignols with *p*-coumarate. Journal of Biological Chemistry 287: 8347–8355.

Wu F, Wu Z, Yang A, Jiang S, Wang Z-Y, Fu C. 2018. Functional characterization of caffeic acid O-methyltransferase in internode lignification of switchgrass (*Panicum virgatum*). Frontiers of Agricultural Science and Engineering 5: 98–107.

Wu Z, Wang N, Hisano H, Cao Y, Wu F, Liu W, Bao Y, Wang ZY, Fu C. 2019. Simultaneous regulation of *F5H* in COMT-RNAi transgenic switchgrass alters effects of *COMT* suppression on syringyl lignin biosynthesis. Plant Biotechnology Journal 17: 836–845.

Yamamoto S, Afifi OA, Lam LPY, Takeda-Kimura Y, Osakabe Y, Osakabe K, Bartley LE, Umezawa T, Tobimatsu Y. 2024. Disruption of aldehyde dehydrogenase decreases cell wall-bound *p*-hydroxycinnamates and improves cell wall digestibility in rice. The Plant Journal 120: 2828–2845.

Yamamura M, Wada S, Sakakibara N, Nakatsubo T, Suzuki S, Hattori T, Takeda M, Sakurai N, Suzuki H, Shibata D, et al. 2011. Occurrence of guaiacyl/*p*-hydroxyphenyl lignin in *Arabidopsis thaliana* T87 cells. Plant Biotechnology 28: 1–8.

