## Supporting Information for "Stacked mutations disrupting syringyl and *p*-coumaroylated lignin biosynthesis in rice result in lignin dominated by guaiacyl units: insights into grass-specific lignin monomer biosynthesis and polymerization mechanisms"

**List of Materials**

**Fig. S1.** DFRC-derived lignin monomers released from rice culm cell walls.

**Fig. S2.** 2D HSQC NMR spectra of whole rice culm cell walls.

**Fig. S3.** Volume integration analysis of 2D HSQC NMR spectra of whole rice culm cell walls.

**Fig. S4.** 2D HSQC NMR spectra of dioxane/water-soluble lignins extracted from rice mutants.

**Fig. S5.** NMR-based lignin linkage distribution analysis of dioxane/water-soluble lignins.

**Table S1.** Primers and oligonucleotides used in this study.

**Table S2.** Accession numbers of *CAld5H* and *PMT* genes studied.

**Table S3.** Growth characteristics of rice mutant lines.

**Table S4.** Yield and composition of DFRC-derived lignin monomers.

**Table S5.** Peak assignments in 2D HSQC NMR spectra of rice cell walls.

**Table S6.** Peak assignments in 2D HSQC NMR spectra of dioxane/water-soluble lignins.

**Table S7.** Molecular weight distribution data of dioxane/water-soluble lignins and synthetic lignin.

**Supplementary References**

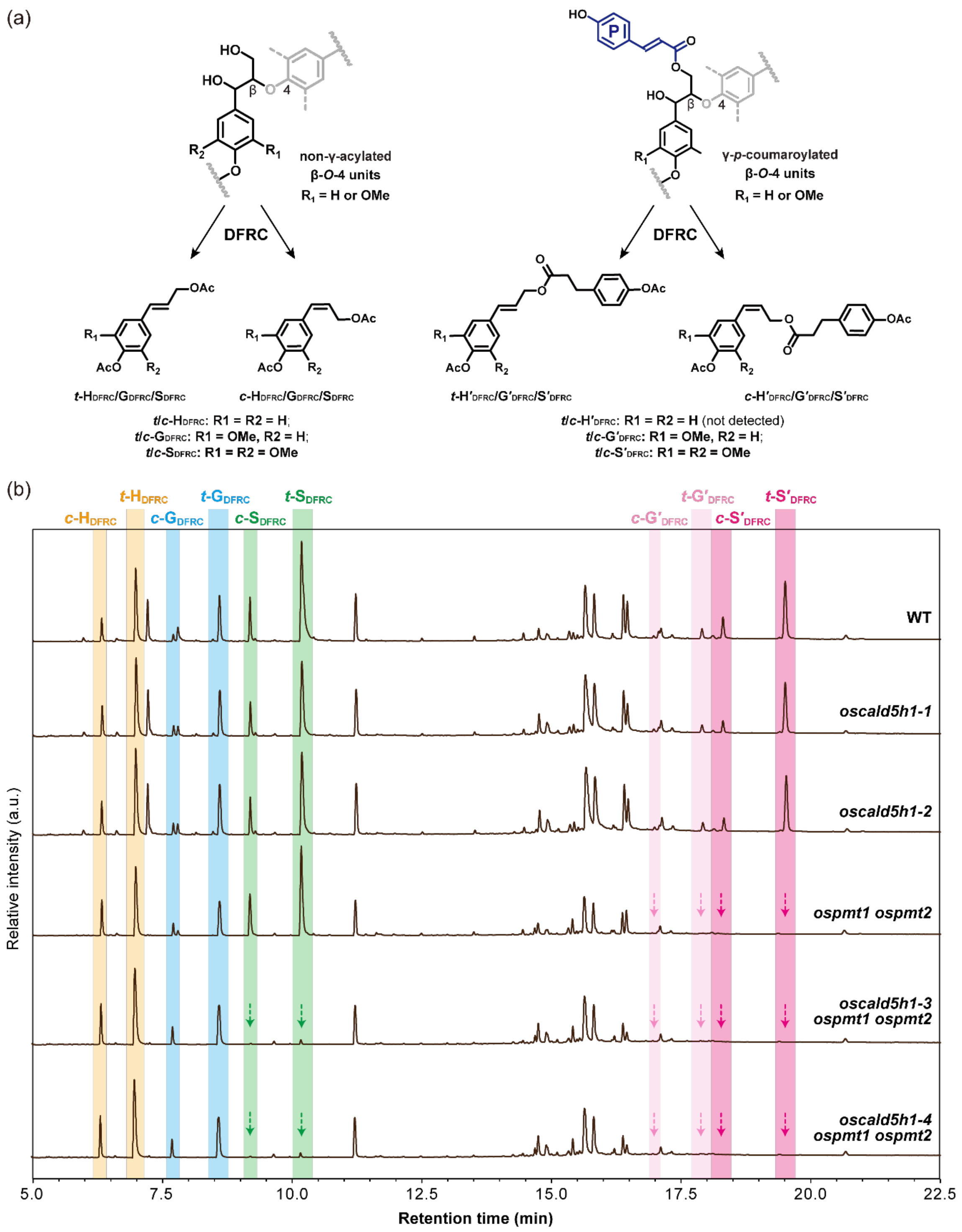


**Fig. S1. DFRC-derived lignin monomers released from rice culm cell walls.** (a) Non-γ-acylated (**H**$_{\mathbf{DFRC}}$, **G**$_{\mathbf{DFRC}}$ and **S**$_{\mathbf{DFRC}}$) and γ-*p*-coumaroylated (**G′**$_{\mathbf{DFRC}}$ and **S′**$_{\mathbf{DFRC}}$) lignin degradation monomers released by cleavages of non-γ-acylated and γ-*p*-coumaroylated β–O–4 units via derivatization followed by reductive cleavage (DFRC) reaction. (b) GC-MS chromatograms of DFRC-derived lignin monomers released from rice culm cell walls. The peaks from *trans*- and *cis*-isomers of DFRC-derived lignin degradation monomers are indicated. WT, wild type line; *oscald5h1-1* and *oscald5h1-2*, *OsCAld5H1* single-knockout lines; *ospmt1 ospmt2*, *OsPMT1* and *OsPMT2* double-knockout lines; *oscald5h1-3 ospmt1 ospmt2* and *oscald5h1-4 ospmt1 ospmt2*, *OsCAld5H1*, *OsPMT1* and *OsPMT2* triple-knockout lines.

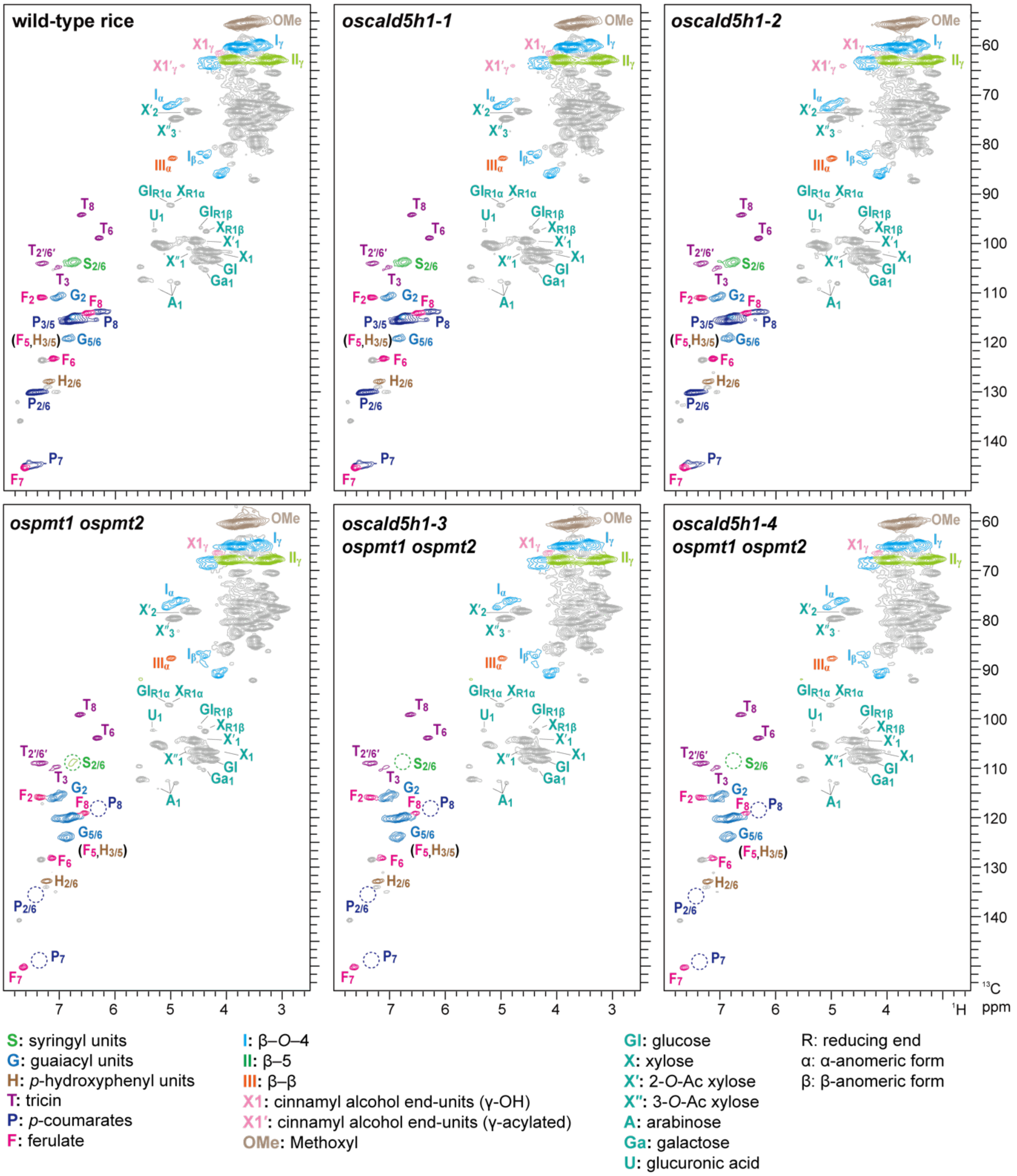


**Fig. S2. 2D HSQC NMR spectra of whole rice culm cell walls.** Ball-milled cell wall residue (CWR) samples from mature rice culms were subjected to whole cell wall NMR analysis using the dimeyhylsulfoxide-$d_6$/pyridine-$d_5$ solvent system (Kim and Ralph, 2010; Mansfield et al., 2012). Volume integration data and signal assignments are listed in **Supplementary Figure S3** and **Supplementary Table S5**. WT, wild-type line; *oscald5h1-1* and *oscald5h1-2*, *OsCAld5H1* single-knockout lines; *ospmt1 ospmt2*, *OsPMT1* and *OsPMT2* double-knockout lines; *oscald5h1-3 ospmt1 ospmt2* and *oscald5h1-4 ospmt1 ospmt2*, *OsCAld5H1*, *OsPMT1* and *OsPMT2* triple-knockout lines.

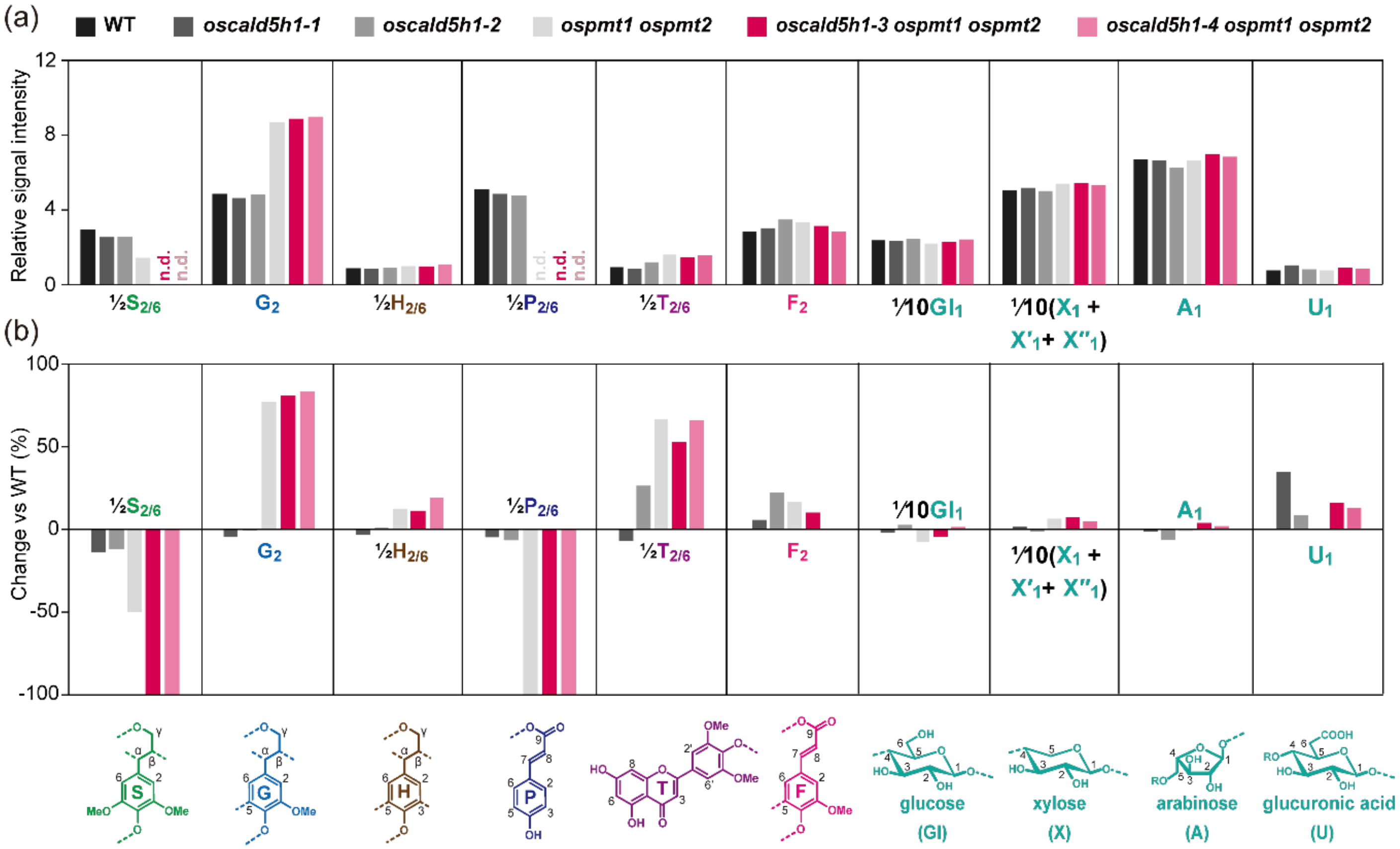


**Fig. S3. Volume integration analysis of 2D HSQC NMR spectra of whole rice culm cell walls.** (a) Normalized signal intensities of major lignin, hydroxycinnamate, and polysaccharide units expressed as percentages of the sum of the listed signals (½$\mathbf{S_{2/6}}$ + $\mathbf{G_2}$ + ½$\mathbf{P_{2/6}}$ + $\mathbf{F_2}$ + ½$\mathbf{T_{2'/6'}}$ + $\mathbf{A_1}$ + $\mathbf{U_1}$ + $\mathbf{Gl_1}$ + $\mathbf{X_1}$ + $\mathbf{X'_1}$ + $\mathbf{X''_1}$ = 100). (b) The extent of changes in lignin, hydroxycinnamate, and polysaccharide signal intensities in mutants compared to the wild-type control. n.d., not detected; S, syringyl units; G, guaiacyl units; H, *p*-hydroxyphenyl units; P, *p*-coumarate units; T, tricin units; F, ferulate units; Gl, glucose; X, xylose; A, arabinose; U, glucuronic acid; WT, wild type line; *oscald5h1-1* and *oscald5h1-2*, *OsCAld5H1* single-knockout lines; *ospmt1 ospmt2*, *OsPMT1* and *OsPMT2* double-knockout lines; *oscald5h1-3 ospmt1 ospmt2* and *oscald5h1-4 ospmt1 ospmt2*, *OsCAld5H1*, *OsPMT1* and *OsPMT2* triple-knockout lines.

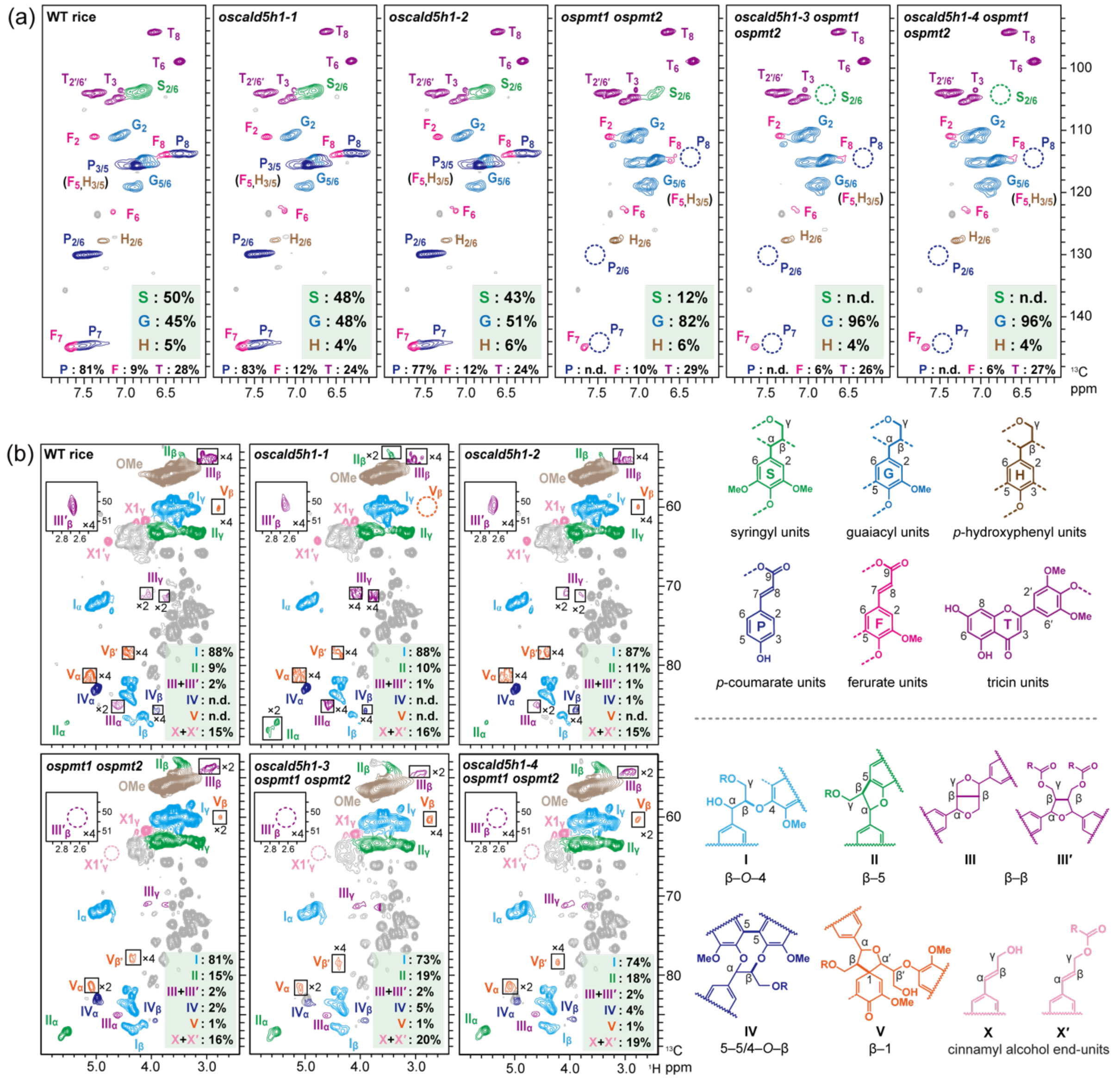


**Fig. S4. 2D HSQC NMR spectra of dioxane/water-soluble lignins extracted from rice mutants.** Aromatic (a) and oxygenated aliphatic (b) sub-regions of 2D HSQC NMR spectra are displayed. The soluble lignin fractions were extracted by dioxane-water (96:4 v/v) from rice culm cell walls. Contour coloration matches the substructures shown in each panel. Peak assignments are listed in **Supplementary Table S6**. Volume integrals are given for the major lignin aromatic units as percentages relative to the total of syringyl (**S**), guaiacyl (**G**), *p*-hydroxyphenyl (**H**) aromatic units (½$\mathbf{S_{2/6}}$ + $\mathbf{G_2}$ + ½$\mathbf{H_{2/6}}$ = 100%) in (a) and for the major lignin inter-monomeric linkage and end-unit types as percentages relative to the total of the inter-monomeric linkage types ($\mathbf{I_\alpha}$ + $\mathbf{II_\alpha}$ + ½$\mathbf{III_\alpha}$ + ½$\mathbf{III'_\beta}$ + $\mathbf{IV_\alpha}$ + $\mathbf{V_\alpha}$ = 100%) in (b). WT, wild type line; *oscald5h1-1* and *oscald5h1-2*, *OsCAld5H1* single-knockout lines; *ospmt1 ospmt2*, *OsPMT1* and *OsPMT2* double-knockout lines; *oscald5h1-3 ospmt1 ospmt2* and *oscald5h1-4 ospmt1 ospmt2*, *OsCAld5H1*, *OsPMT1* and *OsPMT2* triple-knockout lines.

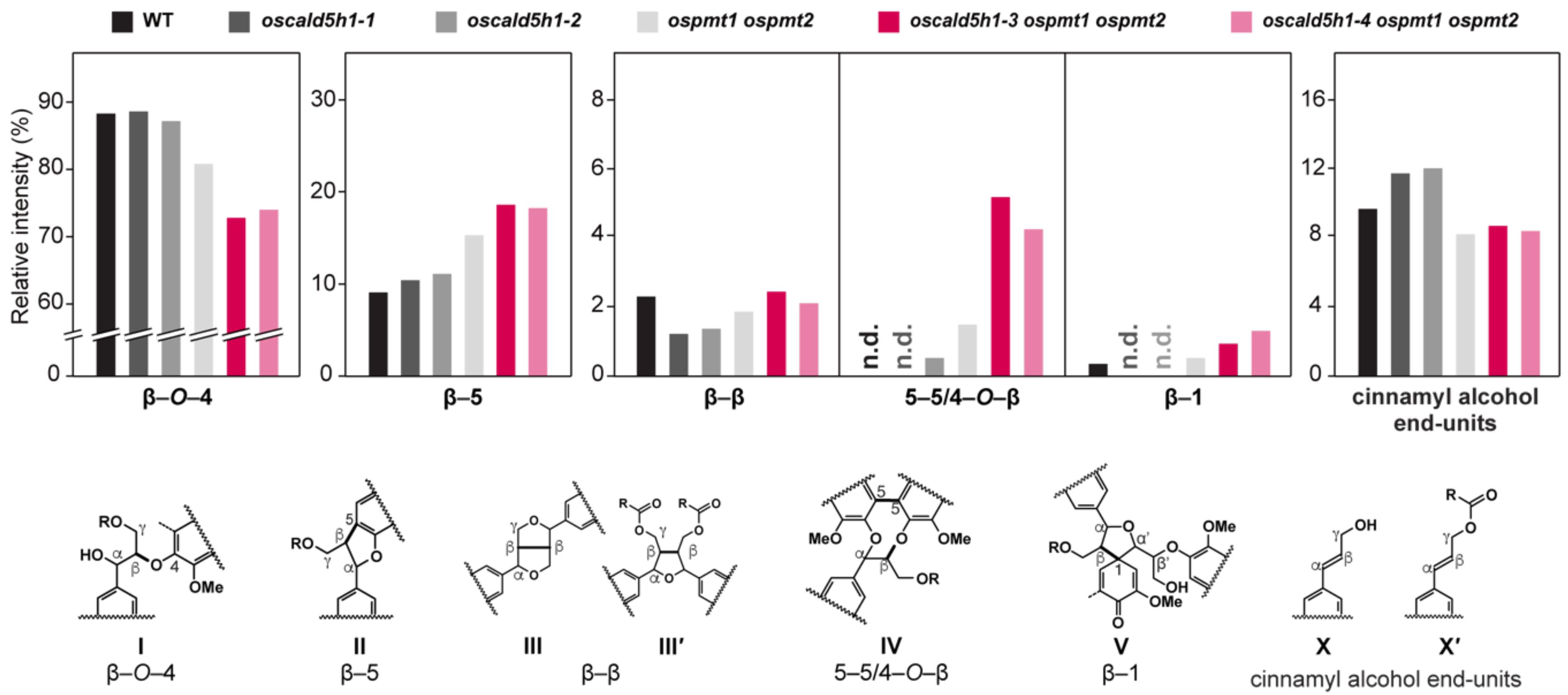


**Fig. S5. NMR-based lignin linkage distribution analysis of dioxane/water-soluble lignins from rice mutants.** Comparisons of volume integration data of lignin linkage types between dioxane/water-soluble lignin fractions extracted from rice wild-type and mutant lines. The HSQC NMR spectra of the dioxane/water-soluble lignin samples and peak assignments are shown in **Supplementary Figure S4** and **Supplementary Table S6**. Volume integral analysis was performed for the major lignin inter-monomeric linkage and end-unit types as percentages relative to the total of the analyzed inter-monomeric linkage types ($\mathbf{I}_{\alpha}$ + $\mathbf{II}_{\alpha}$ + ½$\mathbf{III}_{\alpha}$ + $\mathbf{IV}_{\alpha}$ + $\mathbf{V}_{\alpha}$ = 100%). WT, wild type line; *oscald5h1-1* and *oscald5h1-2*, *OsCAld5H1* single-knockout lines; *ospmt1 ospmt2*, *OsPMT1* and *OsPMT2* double-knockout lines; *oscald5h1-3 ospmt1 ospmt2* and *oscald5h1-4 ospmt1 ospmt2*, *OsCAld5H1*, *OsPMT1* and *OsPMT2* triple-knockout lines.

**Table S1. Primers and oligonucleotides used in this study.**

| Purpose | Sequence (from 5´ to 3´) |
|---|---|
| Construction of sgRNA-Cas9 expression vectors for generating *oscald5h1 ospmt1 ospmt2* | TTGGGTCTCGTGCATCGCCAGCGCTGCCAGGCCGGTTTTAGAGCTAGAAATAGCA<br>TTGGGTCTCCAAACCGCGGTCGTAGGTGAGGTAGTGCACCAGCCGGGAATCGAA |
| Genotyping of *OsCAld5H1 allele in oscald5h1-3 ospmt1 ospmt2* and *oscald5h1-4 ospmt1 ospmt2* | ATGGCGGACATGGTGAAGTT<br>ACGCACAGCTTCCTCATCTG |
| Genotyping of *OsCAld5H1 allele in oscald5h1-1* and *oscald5h1-2* | GAGCGTGTTCTGCAGGTCGT<br>GATCGTCGGCAACATGGCGA |
| Genotyping of *OsPMT1 allele in ospmt1 ospmt2*, *oscald5h1-3 ospmt1 ospmt2* and *oscald5h1-4 ospmt1 ospmt2* | CTGGAGCTGTCCATCATCG<br>CGTGTGGAAAATGAGGGAGT |
| Genotyping of *OsPMT2 allele in ospmt1 ospmt2*, *oscald5h1-3 ospmt1 ospmt2* and *oscald5h1-4 ospmt1 ospmt2* | GATTCACGGTGACGAGGAC<br>AGCAATATGATCATCGATCCAG |

**Table S2. Accession numbers of *CAld5H* and *PMT* genes studied.**

| Gene name | Alternative name | Gene locus | Protein accession | Reference |
|---|---|---|---|---|
| *OsCAld5H1* | *OsF5H1* | LOC_Os10g36848 | AK067847 | Takeda et al., 2017<br>Takeda et al., 2019 |
| *AtCAld5H1* | *AtF5H1/AtFAH1* | AT4g36220 | Q42600 | Chapple et al., 1992 |
| *OsPMT1* | *OsAT4* | LOC_Os01g18744 | AK060689 | Withers et al., 2012<br>Lam et al., 2024 |
| *OsPMT2* | *OsAT3* | LOC_Os05g04584 | AK069868 | Lam et al., 2024 |

**Table S3. Growth characteristics of rice wild-type and mutants.**

| Trait | WT | *oscald5h1-1* | *oscald5h1-2* | *ospmt1 ospmt2* | *oscald5h1-3 ospmt1 ospmt2* | *oscald5h1-4 ospmt1 ospmt2* |
|---|---|---|---|---|---|---|
| Plant height (cm) [1] | 118.1 ± 2.0 [a] | **102.5 ± 4.0** [b] | **103.2 ± 3.0** [b] | **98.8 ± 3.4** [b] | **91.3 ± 4.2** [c] | **92.3 ± 4.5** [c] |
| Culm length (cm) [2] | 76.3 ± 3.1 [a] | **67.0 ± 1.4** [b] | **65.0 ± 3.2** [b] | **56.7 ± 4.1** [c] | **49.5 ± 3.3** [d] | **52.3 ± 3.0** [cd] |
| Tiller number | 6.0 ± 1.2 [a] | 5.8 ± 1.3 [ab] | 5.0 ± 1.0 [ab] | 5.0 ± 0.0 [ab] | **4.0 ± 1.1** [b] | 4.5 ± 0.8 [ab] |
| Culm CWR yield (%) | 56.9 ± 3.5 [b] | 62.5 ± 3.0 [ab] | 60.4 ± 3.2 [ab] | 62.0 ± 2.4 [ab] | **63.0 ± 2.4** [a] | 61.8 ± 3.1 [ab] |
| Panicle length(cm) | 16.1 ± 0.6 [a] | **13.3 ± 1.3** [b] | 14.5 ± 0.5 [ab] | **11.5 ± 0.5** [b] | **11.5 ± 1.0** [b] | **13.0 ± 1.1** [b] |
| Panicle number | 6.0 ± 1.2 [a] | 5.8 ± 1.3 [ab] | 5.0 ± 1.0 [ab] | 5.0 ± 0.0 [ab] | **4.0 ± 1.1** [b] | 4.5 ± 0.8 [ab] |

[1]Length measured from the soil surface to the tip of the top leaf. [2]Length measured from soil surface to the panicle base. [3]Dry weight of all aerial parts excluding panicles.Values are means ± standard deviation from biological replicates (n ≥ 4). Different letters on values indicate significant differences (one-way ANOVA with Tukey's test, $P < 0.05$). Bold values indicate statistically significant compared to the WT control. WT, wild type line; *oscald5h1-1* and *oscald5h1-2*, *OsCAld5H1* single-knockout lines; *ospmt1 ospmt2*, *OsPMT1* and *OsPMT2* double-knockout lines; *oscald5h1-3 ospmt1 ospmt2* and *oscald5h1-4 ospmt1 ospmt2*, *OsCAld5H1*, *OsPMT1* and *OsPMT2* triple-knockout lines.

**Table S4. Yield and composition of DFRC-derived lignin monomers released from rice culm cell walls.**

| | WT | *oscald5h1-1* | *oscald5h1-2* | *ospmt1 ospmt2* | *oscald5h1-3 ospmt1 ospmt2* | *oscald5h1-4 ospmt1 ospmt2* |
|---|---|---|---|---|---|---|
| *Monomer Yield (μmol/g CWR)* | | | | | | |
| **$H_{DFRC}$** | 4.21 ± 0.09 [cd] | 4.00 ± 0.19 [d] | 4.47 ± 0.17 [c] | **5.90 ± 0.10 [a]** | **5.03 ± 0.08 [b]** | **6.13 ± 0.20 [a]** |
| **$G_{DFRC}$** | 18.90 ± 0.73 [b] | 18.23 ± 0.74 [b] | 19.33 ± 1.28 [b] | **24.68 ± 0.34 [a]** | **24.59 ± 1.59 [a]** | **27.18 ± 0.69 [a]** |
| **$S_{DFRC}$** | 10.50 ± 0.45 [a] | **4.16 ± 2.04 [b]** | **5.62 ± 0.42 [b]** | 9.65 ± 0.20 [a] | **trace** | **trace** |
| **$G'_{DFRC}$** | 4.01 ± 0.45 [a] | 3.06 ± 1.07 [ab] | **2.54 ± 0.23 [b]** | **n.d.** | **n.d.** | **n.d.** |
| **$S'_{DFRC}$** | 17.55 ± 1.11 [a] | **10.78 ± 4.18 [b]** | **11.66 ± 1.27 [b]** | **n.d.** | **n.d.** | **n.d.** |
| **Total** | 55.16 ± 2.45 [a] | **40.23 ± 7.16 [bc]** | **43.62 ± 1.01 [b]** | **40.23 ± 0.61 [bc]** | **29.92 ± 1.68 [c]** | **33.56 ± 0.75 [c]** |
| *Monomer Composition (%)* | | | | | | |
| **%$H_{DFRC}$** | 7.63 ± 0.05 [d] | **10.15 ± 1.66 [c]** | **10.24 ± 0.20 [c]** | **14.66 ± 0.16 [b]** | **16.84 ± 0.70 [a]** | **18.28 ± 0.59 [a]** |
| **%$G_{DFRC}$** | 34.27 ± 0.01 [d] | **46.31 ± 8.50 [c]** | **44.31 ± 2.47 [c]** | **61.36 ± 0.22 [b]** | **82.15 ± 0.73 [a]** | **81.00 ± 0.59 [a]** |
| **%$S_{DFRC}$** | 19.05 ± 0.19 [b] | **9.94 ± 3.77 [c]** | **12.88 ± 0.82 [c]** | **23.98 ± 0.14 [a]** | **n.d.** | **n.d.** |
| **%$G'_{DFRC}$** | 7.25 ± 0.83 [a] | 7.51 ± 1.58 [a] | 5.82 ± 0.50 [a] | **n.d.** | **n.d.** | **n.d.** |
| **%$S'_{DFRC}$** | 31.80 ± 0.62 [a] | 26.10 ± 5.98 [a] | 26.75 ± 3.01 [a] | **n.d.** | **n.d.** | **n.d.** |

Values are means ± standard deviation of biological replicates ($n$ = 3). Different letters on values indicate significant differences (one-way ANOVA with Tukey's test, $P < 0.05$). Bold values indicate statistically significant compared to the WT control. CWR, cell wall residue; n.d., not detected; WT, wild type line; *oscald5h1-1* and *oscald5h1-2*, *OsCAld5H1* single-knockout lines; *ospmt1 ospmt2*, *OsPMT1* and *OsPMT2* double-knockout lines; *oscald5h1-3 ospmt1 ospmt2* and *oscald5h1-4 ospmt1 ospmt2*, *OsCAld5H1*, *OsPMT1* and *OsPMT2* triple-knockout lines.

**Table S5. Peak assignments in 2D HSQC NMR spectra of rice cell walls.**

| Labels | $\delta_C/\delta_H$ (ppm) | Assignment |
|---|---|---|
| *Lignin and hydroxycinnamate signals* | | |
| **$S_{2/6}$** | 103.92/6.743 | $C_2$–$H_2$ and $C_6$–$H_6$ in syringyl units |
| **$G_2$** | 110.97/7.034 | $C_2$–$H_2$ in guaiacyl units |
| **$G_{5/6}$** | 119.26/6.839, 115.82/6.826 | $C_5$–$H_5$ and $C_6$–$H_6$ in guaiacyl units |
| **$H_{2/6}$** | 127.98/7.184 | $C_2$–$H_2$ and $C_6$–$H_6$ in *p*-hydroxyphenyl units |
| **$H_{3/5}$** | 115.82/6.826 | $C_3$–$H_3$ and $C_5$–$H_5$ in *p*-hydroxyphenyl units |
| **$T_3$** | 104.89/7.036 | $C_3$–$H_3$ in tricin residues |
| **$T_6$** | 99.07/6.298 | $C_6$–$H_6$ in tricin residues |
| **$T_8$** | 94.4/6.616 | $C_8$–$H_8$ in tricin residues |
| **$T_{2'/6'}$** | 104.18/7.31 | $C_{2'}$–$H_{2'}$ and $C_{6'}$–$H_{6'}$ in tricin residues |
| **$P_{2/6}$** | 130.18/7.449 | $C_2$–$H_2$ and $C_6$–$H_6$ in *p*-coumarate residues |
| **$P_{3/5}$** | 115.82/6.826 | $C_3$–$H_3$ and $C_5$–$H_5$ in *p*-coumarate residues |
| **$P_7$** | 144.9/7.469 | $C_7$–$H_7$ in *p*-coumarate residues |
| **$P_8$** | 113.88/6.278 | $C_8$–$H_8$ in *p*-coumarate residues |
| **$F_2$** | 111.06/7.328 | $C_2$–$H_2$ in ferulate residues |
| **$F_5$** | 115.82/6.826 | $C_5$–$H_5$ in ferulate residues |
| **$F_6$** | 123.31/7.113 | $C_6$–$H_6$ in ferulate residues |
| **$F_7$** | 145.43/7.625 | $C_7$–$H_7$ in ferulate residues |
| **$F_8$** | 114.23/6.517 | $C_8$–$H_8$ in ferulate residues |
| **$I_\alpha$** | 72.72/5.063 | $C_\alpha$–$H_\alpha$ in β–*O*–4 units |
| **$I_\beta$** | 82.64/4.367, 86.57/4.123 | $C_\beta$–$C_\beta$ in β–*O*–4 units |
| **$I_\gamma$** | 60.48/3.75 | $C_\gamma$–$H_\gamma$ in β–*O*–4 units |
| **$II_\gamma$** | 63.43/3.956, 63.34/3.241 | $C_\gamma$–$H_\gamma$ in β–5 substructures |
| **$III_\alpha$** | 82.99/4.974 | $C_\alpha$–$H_\alpha$ in β–β substructures |
| **$X1_\gamma$** | 61.73/4.129 | $C_\gamma$–$H_\gamma$ in γ-free cinnamyl alcohol end-units |
| **$X1'_\gamma$** | 64.32/4.794 | $C_\gamma$–$H_\gamma$ in γ-acylated cinnamyl alcohol end-units |
| **OMe** | 55.66/3.687 | C–H in aromatic methoxyl groups |
| *Polysaccharide anomeric signals* | | |
| **$Gl_1$** | 103.45/4.246 | C1–H1 in (1→4)-β-D-glucopyranosyl units |
| **$X_1$** | 101.97/4.343 | C1–H1 in (1→4)-β-D-xylopyranosyl units |
| **$X'_1$** | 99.65/4.569 | C1–H1 in 2-*O*-acetyl-β-D-xylopyranosyl units |
| **$X''_1$** | 101.14/4.682 | C1–H1 in 3-*O*-acetyl-β-D-xylopyranosyl units |
| **$A_1$** | 109.38/5.216, 108.08/4.881, 107.71/4.997 | C1–H1 in α-L-arabinofuranosyl units |
| **$Ga_1$** | 105.49/4.386 | C1–H1 in (1→4)-β-D-galactopyranosyl units |
| **$U_1$** | 97.52/5.301 | C1–H1 in 4-*O*-methyl-α-D-glucuronopyranosyl units |

Measured in DMSO-$d_6$/pyridine-$d_5$ (4:1, v/v). Signal assignment was based on comparison with NMR data in literature (Kim and Ralph, 2010; Mansfield et al., 2012; Lan et al., 2015; 2018; Afifi et al., 2022; Martin et al., 2023; Lam et al., 2024; Ralph et al. 2024).

**Table S6. Peak assignments in 2D HSQC NMR spectra of dioxane/water-soluble lignins.**

| Labels | $\delta_C/\delta_H$ (ppm) | Assignment |
|---|---|---|
| **$S_{2/6}$** | 103.93/6.782 | $C_2$–$H_2$ and $C_6$–$H_6$ in syringyl units |
| **$G_2$** | 111.1/7.061 | $C_2$–$H_2$ in guaiacyl units |
| **$G_{5/6}$** | 119.38/6.866, 115.79/6.84 | $C_5$–$H_5$ and $C_6$–$H_6$ in guaiacyl units |
| **$H_{2/6}$** | 127.98/7.268 | $C_2$–$H_2$ and $C_6$–$H_6$ in *p*-hydroxyphenyl units |
| **$H_{3/5}$** | 115.79/6.843 | $C_3$–$H_3$ and $C_5$–$H_5$ in *p*-hydroxyphenyl units |
| **$T_3$** | 104.96/7.071 | $C_3$–$H_3$ in tricin residues |
| **$T_6$** | 99.07/6.32 | $C_6$–$H_6$ in tricin residues |
| **$T_8$** | 94.37/6.627 | $C_8$–$H_8$ in tricin residues |
| **$T_{2'/6'}$** | 104.33/7.364 | $C_{2'}$–$H_{2'}$ and $C_{6'}$–$H_{6'}$ in tricin residues |
| **$P_{2/6}$** | 130.21/7.466 | $C_2$–$H_2$ and $C_6$–$H_6$ in *p*-coumarate residues |
| **$P_{3/5}$** | 115.79/6.843 | $C_3$–$H_3$ and $C_5$–$H_5$ in *p*-coumarate residues |
| **$P_7$** | 144.78/7.504 | $C_7$–$H_7$ in *p*-coumarate residues |
| **$P_8$** | 113.88/6.323 | $C_8$–$H_8$ in *p*-coumarate residues |
| **$F_2$** | 111.1/7.366 | $C_2$–$H_2$ in ferulate residues |
| **$F_5$** | 115.79/6.843 | $C_5$–$H_5$ in ferulate residues |
| **$F_6$** | 123.36/7.134 | $C_6$–$H_6$ in ferulate residues |
| **$F_7$** | 145.02/7.653 | $C_7$–$H_7$ in ferulate residues |
| **$F_8$** | 113.96/6.466 | $C_8$–$H_8$ in ferulate residues |
| **$I_\alpha$** | 72.01/5.017 | $C_\alpha$–$H_\alpha$ in β–*O*–4 units |
| **$I_\beta$** | 83.76/4.507, 86.22/4.209 | $C_\beta$–$C_\beta$ in β–*O*–4 units |
| **$I_\gamma$** | 60.06/3.715 | $C_\gamma$–$H_\gamma$ in β–*O*–4 units |
| **$II_\alpha$** | 87.2/5.527 | $C_\alpha$–$H_\alpha$ in β–5 substructures |
| **$II_\beta$** | 53.31/3.508 | $C_\beta$–$H_\beta$ in β–5 substructures |
| **$II_\gamma$** | 62.87/3.769, 63.29/3.259 | $C_\gamma$–$H_\gamma$ in β–5 substructures |
| **$III_\alpha$** | 85.09/4.657 | $C_\alpha$–$H_\alpha$ in resinol-type β–β substructures |
| **$III_\beta$** | 53.8/2.965 | $C_\beta$–$H_\beta$ in resinol-type β–β substructures |
| **$III_\gamma$** | 70.96/4.126, 71.17/3.777 | $C_\gamma$–$H_\gamma$ in resinol-type β–β substructures |
| **$III'_\beta$** | 50.28/2.67 | $C_\beta$–$H_\beta$ in in tetrahydrofuran-type β–β substructures |
| **$IV_\alpha$** | 82.98/5.009 | $C_\alpha$–$H_\alpha$ in 5–5 substructures |
| **$IV_\beta$** | 85.66/3.972 | $C_\beta$–$C_\beta$ in 5–5 substructures |
| **$V_\alpha$** | 81.44/5.117 | $C_\alpha$–$H_\alpha$ in β–1 substructures |
| **$V_\beta$** | 59.85/2.843 | $C_\beta$–$C_\beta$ in β–1 substructures |
| **$V_{\beta'}$** | 78.48/4.437 | $C_{\beta'}$–$C_{\beta'}$ in β–1 substructures |
| **$X1_\gamma$** | 61.68/4.16 | $C_\gamma$–$H_\gamma$ in γ-free cinnamyl alcohol end-units |
| **$X1'_\gamma$** | 64.28/4.823 | $C_\gamma$–$H_\gamma$ in γ-acylated cinnamyl alcohol end-units |
| **OMe** | 55.7/3.703 | C–H in aromatic methoxyl groups |

Measured in DMSO-*d₆*/pyridine-*d₅* (4:1, v/v). Signal assignment was based on comparison with NMR data in literature (Kim and Ralph, 2010; Mansfield et al., 2012; Lan et al., 2015; 2018; Afifi et al., 2022; Martin et al., 2023; Lam et al., 2024; Ralph et al. 2024).

**Table S7. Molecular weight data of G-dominated lignins from rice, Arabidopsis and pine, and synthetic lignin.**

| | Origin of lignins | $M_n$ | $M_w$ | PDI |
|---|---|---|---|---|
| Rice | Wild-type rice (cv. Nipponbare) | 2900 | 9100 | 3.1 |
| | *oscald5h1-1* | 3000 | 9000 | 3.0 |
| | *oscald5h1-2* | 3000 | 9200 | 3.1 |
| | *ospmt1 ospmt2* | 3000 | 8900 | 3.0 |
| | *oscald5h1-3 ospmt1 ospmt2* | 3000 | 8500 | 2.9 |
| | *oscald5h1-4 ospmt1 ospmt2* | 2900 | 8500 | 2.9 |
| Arabidopsis | Wild-type Arabidopsis (Col-0) | 3700 | 14500 | 4.0 |
| | *atcald5h1* | 3600 | 14900 | 4.0 |
| Pine | *Pinus taeda* | 3100 | 12300 | 3.9 |
| *In vitro* | G-DHP | 2500 | 12200 | 4.8 |

Molecular weight distribution data were obtained by gel permeation chromatography (GPC) using polystyrene molecular weight standards for dioxane/water-soluble lignin (DL) samples extracted from G-lignin-dominated rice (*oscald5h1-3 ospmt1 ospmt2* and *oscald5h1-4 ospmt1 ospmt2*) and Arabidopsis (*atcald5h1*) mutant lines, wild-type rice, Arabidopsis, pine, and synthetic G lignin (G-DHP) prepared by *in vitro* polymerization of coniferyl alcohol. $M_n$, number-averaged molecular weight; $M_w$, weight-averaged molecular weight; PDI, polydiversity index ($M_w/M_n$).

### Supplementary References


**Afifi OA, Tobimatsu Y, Lam PY, Martin AF, Miyamoto T, Osakabe Y, Osakabe K, Umezawa T. 2022.** Genome-edited rice deficient in two *4-COUMARATE:COENZYME A LIGASE* genes displays diverse lignin alterations. *Plant Physiology* **190**: 2155–2172.

**Chapple C, Vogt T, Ellis BE, Somerville CR. 1992.** An Arabidopsis mutant defective in the general phenylpropanoid pathway. *Plant Cell*, **4:** 1413−1424.

**Kim H, Ralph J. 2010.** Solution-state 2D NMR of ball-milled plant cell wall gels in DMSO-$d_6$/pyridine-$d_5$. *Organic & Biomololecular Chemistry* **8:** 576–591.

**Lam LPY, Tobimatsu Y, Suzuki S, Tanaka T, Yamamoto S, Takeda-Kimura Y, Osakabe Y, Osakabe K, Ralph J, Bartley LE, Umezawa T. 2024.** Disruption of *p*-coumaroyl-CoA:monolignol transferases in rice drastically alters lignin composition. *Plant Physiology*, **194:** 832−848.

**Lan W, Lu F, Regner M, Zhu Y, Rencoret J, Ralph SA, Zakai UI, Morreel K, Boerjan W, Ralph J. 2015.** Tricin, a flavonoid monomer in monocot lignification. *Plant Physiology* **167:** 1284-1295.

**Lan W, Yue F, Rencoret J, Del Río JC, Boerjan W, Lu F, Ralph J. 2018.** Elucidating tricin-lignin structures: assigning correlations in HSQC spectra of monocot lignins. *Polymers* **10:** 916.

**Mansfield SD, Kim H, Lu F, Ralph J. 2012.** Whole plant cell wall characterization using solution-state 2D NMR. *Nature Protocols* **7:** 1579–1589.

**Martin AF, Tobimatsu Y, Lam PY, Matsumoto N, Tanaka T, Suzuki S, Kusumi R, Miyamoto T, Takeda-Kimura Y, Yamamura M, Koshiba T, Osakabe K, Osakabe Y, Sakamoto M, Umezawa T.** 2023. Lignocellulose molecular assembly and deconstruction properties of lignin-altered rice mutants. *Plant Physiology* **191:** 70–86.

**Ralph SA, Ralph J, Lu F. 2024.** NMR database of lignin and cell wall model compounds. Available at https://doi.org/10.11578/2409191.

**Takeda Y, Koshiba T, Tobimatsu Y, Suzuki S, Murakami S, Yamamura M, Rahman MM, Takano T, Hattori T, Sakamoto M, Umezawa T. 2017.** Regulation of *CONIFERALDEHYDE 5-HYDROXYLASE* expression to modulate cell wall lignin structure in rice. *Planta*, **246:** 337−349.

**Takeda Y, Suzuki S, Tobimatsu Y, Osakabe K, Osakabe Y, Ragamustari SK, Sakamoto M, Umezawa T. 2019.** Lignin characterization of rice *CONIFERALDEHYDE 5-HYDROXYLASE* loss-of-function mutants generated with the CRISPR/Cas9 system. *The Plant Journal*, **97:** 543−554.

**Withers S, Lu F, Kim H, Zhu Y, Ralph J, Wilkerson CG. 2012.** Identification of grass-specific enzyme that acylates monolignols with *p*-coumarate. *Journal of Biological Chemistry*, **287:** 8347−8355.
